# Golgi-endosomal CATCHR complexes function as organizing hubs for vesicle tethering and fusion

**DOI:** 10.64898/2026.06.16.732723

**Authors:** Walter S. Aragon-Ramirez, Reece C. Forrest, Irina Pokrovskaya, Amrita Khakurel, Farhana Taher Sumya, Vladimir V. Lupashin

## Abstract

Approximately one-third of all human proteins transit through the secretory pathway, where the Golgi apparatus orchestrates protein modification, sorting, and distribution through highly selective vesicle budding and fusion events. Central to these processes are the Complexes Associated with Tethering Containing Helical Rods (CATCHR), multisubunit tethering complexes that coordinate vesicle docking and fusion through interactions with coiled-coil tethers (CCTs), Rab GTPases, SNAREs, and Sec1/Munc18 (SM) proteins and other trafficking factors. To define the molecular organization of endosomal-Golgi CATCHR complexes, we generated the first comprehensive proximity map of the COG, GARP, and EARP tethering complexes using functional, near-endogenously expressed TurboID-tagged subunits. Comparative proximity proteomics revealed that each CATCHR complex assembles a distinct trafficking module composed of characteristic CCTs, Rab-associated proteins, SNAREs, and SM proteins, establishing a system-level framework for the spatial organization of Golgi and endosomal membrane trafficking. The COG complex preferentially associates with Golgi CCTs and the STX5-SCFD1 fusion machinery, GARP with CCDC186 and *Trans*-Golgi Network (TGN) SNAREs, and EARP with GRIPAP1, the VPS33B-VIPAS39 (CHEVI) complex, and RAB11-dependent recycling machinery. Beyond validating known interactions, our study identifies CCDC186 as a vesicle tether, establishes WWOX as a previously unrecognized regulator of Golgi homeostasis and glycosylation, and provides evidence that Golgi CATCHR complexes function as central organizing hubs that assemble specialized trafficking modules to coordinate vesicle tethering and membrane fusion.

## Introduction

The Golgi apparatus is the central organelle in charge of modifying, packaging, and sorting proteins in the secretory pathway. It is estimated that one-third of the human genome encodes proteins whose functions depend on their proper traffic through the Golgi^1^. Golgi trafficking is a finely regulated process that depends on the specific budding, tethering, and fusion of vesicles^2^. Vesicular tethering and fusion is a selective process where multiple proteins participate to deliver cargoes to their proper acceptor intracellular compartment^2^. The Complexes Associated with Tethering Containing Helical Rods (CATCHR) are an evolutionarily conserved family of proteins that have been proposed to play key roles in regulating vesicular tethering and fusion at different stages of the secretory pathway^3,4^. The CATCHR family is composed of five members: NRZ, COG, GARP, EARP, and Exocyst. The NRZ complex is composed of three subunits and has been proposed to facilitate the tethering of vesicles on endoplasmic reticulum membranes^5^. The Exocyst complex is composed of eight subunits, and it has been proposed to tether vesicles at the plasma membrane^6^. The COG and GARP complexes carry out their tethering functions on Golgi membranes, with the COG complex functioning in tethering vesicles at multiple Golgi subcompartments^7,8^ and the GARP complex functioning at the *Trans*-Golgi Network (TGN)^9,10^. The EARP complex has been proposed to work as a tethering complex in the endolysosomal system^11^ (**Figure 1A**).

**Figure 1.**
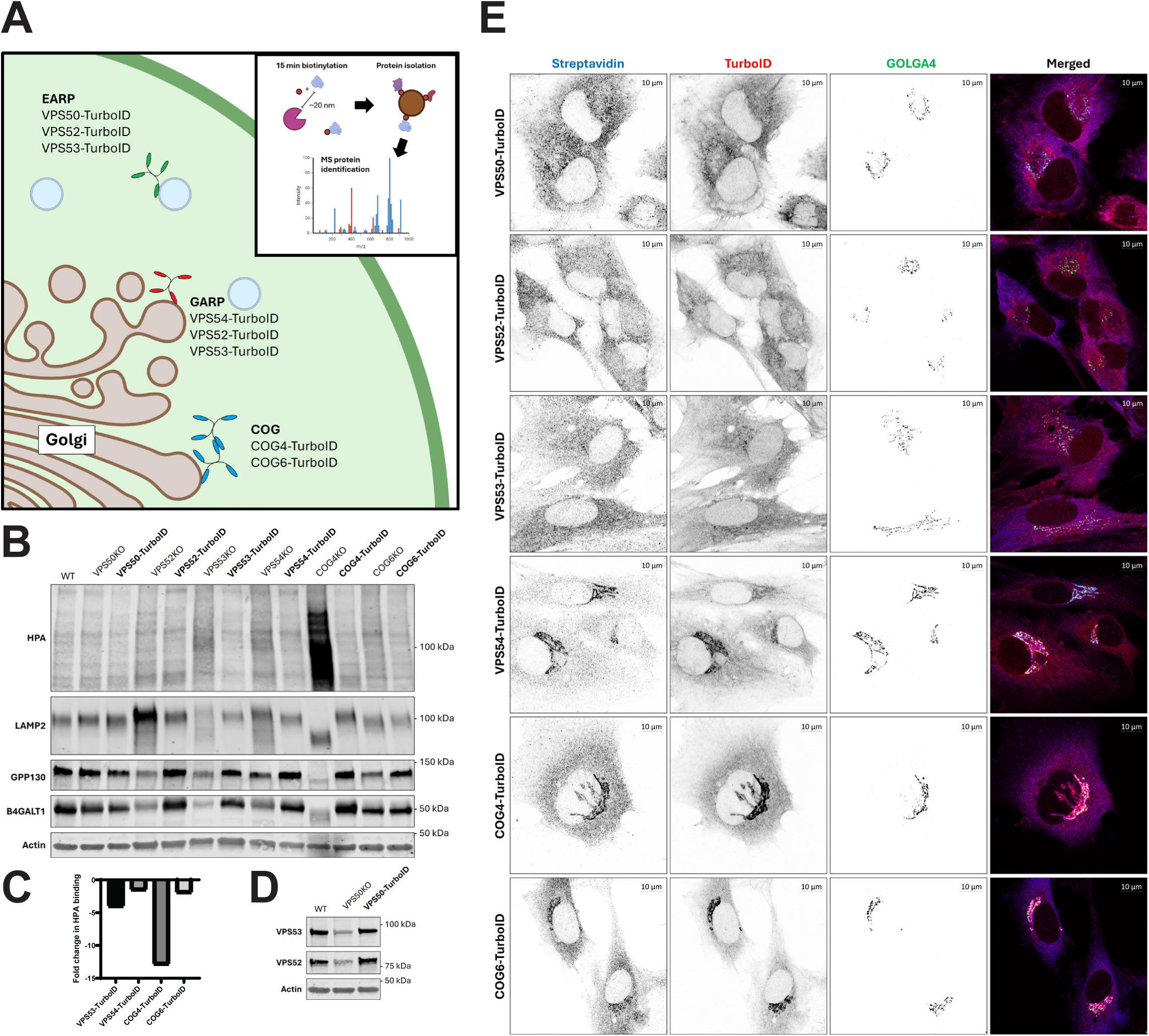
Endogenous TurboID tagging preserves CATCHR complex function and enables compartment-specific proximity labeling. (**A**) Schematic representation of the Golgi and endolysosomal CATCHR complexes analyzed in this study, including EARP (VPS52-, VPS53-, and VPS50-TurboID), GARP (VPS52-, VPS53-, and VPS54-TurboID), and COG (COG4- and COG6-TurboID). TurboID-mediated proximity labeling was performed by biotin supplementation followed by streptavidin purification and mass spectrometry analysis. (**B**) Functional validation of the TurboID cell lines. Near-endogenous expression of CATCHR-TurboID fusion proteins rescues trafficking and glycosylation defects associated with depletion of the corresponding endogenous proteins. Rescue was assessed by restoration of O-glycosylation, measured by HPA lectin binding, normalization of LAMP2 glycosylation and abundance, recovery of GPP130 abundance, and restoration of B4GALT1 levels. Actin serves as a loading control. (**C**) Rescue of glycosylation defects in VPS53KO, VPS54KO, COG4KO, and COG6KO cells. HPA binding was quantified in KO cells reconstituted with the corresponding TurboID-tagged protein and is presented as the fold change decrease in HPA signal relative to the parental KO cell line. Reduced HPA binding following rescue indicates restoration of normal glycosylation. (**D**) Expression of VPS50-TurboID restores the stability of the EARP complex, rescuing the abundance of the VPS52 and VPS53 subunits in VPS50 KO cells. (**E**) Immunofluorescence analysis of biotinylated proteins detected with fluorescent streptavidin (first column, left to right) and TurboID-tagged CATCHRs detected by anti-TurboID antibodies (second column) in RPE1 cells stained with Golgi marker GOLGA4/P230 (third column). VPS50-, VPS52-, and VPS53-TurboID localize to endosomal compartments, VPS54-, COG4-, and COG6-TurboID localize to Golgi membranes. Streptavidin staining closely overlaps with each TurboID construct, demonstrating spatially restricted proximity labeling within the expected intracellular compartments.

CATCHR complexes interact with multiple core components of the membrane trafficking machinery, including small GTPases, Coiled-Coil tethers (CCTs), SNAREs, and Sec1/Munc18 (SM) proteins^12–15^. It is well established that the CATCHR complexes are essential for the accurate delivery of transport carriers to their target membranes. Consistent with this central role, mutations in CATCHR subunits cause lethal disorders in humans, and depletion of a single subunit is embryonically lethal in mammals^16–20^. Likewise, knockout (KO) and knockdown (KD) cellular models of CATCHRs deficiency exhibit profound defects in intracellular trafficking, Golgi organization, and protein glycosylation^21–24^. Together, these observations underscore the fundamental importance of CATCHR complexes for membrane trafficking throughout eukaryotes. Despite their central role in membrane trafficking, it remains unclear how individual CATCHR complexes are integrated into the broader trafficking machinery to confer pathway specificity. Although numerous pairwise interactions have been described, a systems-level view of the molecular networks surrounding CATCHR complexes is lacking. To address this knowledge gap, we used near-endogenously expressed TurboID-tagged subunits to generate the first comprehensive proximity map of the Golgi-associated CATCHR complexes COG and GARP, as well as the endosomal CATCHR complex EARP. We uncovered previously unrecognized *in vivo* proximity proteins and new candidate regulators of membrane trafficking, while also identifying known interaction partners and established trafficking factors associated with each complex with high specificity. These findings provide new insights into the organization of CATCHR-dependent trafficking pathways and reveal distinct conserved molecular modules that cooperate with COG, GARP, and EARP to regulate vesicle tethering and fusion. Our analysis reveals that each Golgi-associated CATCHR complex is embedded within a distinct molecular module that reflects its trafficking specialization. For the EARP complex, this module is centered on the sorting and recycling endosome interface and includes the VPS33B-VIPAS39 SM complex alongside regulators of RAB11-dependent recycling, refining the subcellular context in which EARP operates. For GARP, we identify CCDC186 as a bona fide vesicle tether. Mitochondrial relocalization assays combined with transmission electron microscopy (TEM) imaging demonstrate that CCDC186 is sufficient to ectopically capture a specific subset of transport vesicles, providing direct functional evidence for its tethering activity. Unexpectedly, we identify the tumor suppressor WWOX as a previously unrecognized component of the Golgi trafficking machinery. WWOX localizes to the medial Golgi together with the COG complex, and its depletion disrupts Golgi morphology and both N- and O-linked glycosylation, revealing an essential role in Golgi homeostasis. Additionally, WWOX depletion rescues a subset of Golgi recycling proteins mislocalized by acute COG inactivation, suggesting a possible role of WWOX as a regulator of cargo sorting or vesicle formation.

Collectively, our study establishes the first comprehensive *in vivo* proximity map of the Golgi- and endosomal-associated CATCHR complexes COG, GARP, and EARP. Beyond validating known interactions, it defines the molecular architecture of CATCHR-dependent trafficking pathways, uncovers previously unrecognized regulators of Golgi and endosomal transport, and provides a conceptual framework for understanding how distinct tethering complexes achieve pathway specificity during vesicle tethering and fusion.

## Results

### The CATCHR-TurboID constructs are functional and label their proposed compartments and partners

To explore the landscape of the Golgi CATCHRs, we used the TurboID proximity labeling system. TurboID is a fast proximity labeling enzyme engineered from the promiscuous mutant of the bacterial biotin ligase BirA^25^. This system allows the detection of specific transient and stable interactions as it covalently labels (biotinylates) any protein with surface-exposed lysine residues within a ∼10 nm radius of the TurboID-tagged protein. Additionally, this system requires short labeling times, reducing background noise. To decrease the probability of detecting artifacts, we stably expressed C-terminally tagged TurboID CATCHR proteins into their corresponding KO hTERT-RPE1 cell line. We created six cell lines: COG4-TurboID and COG6-TurboID to account for the two lobes of the COG complex; VPS50-TurboID and VPS54-TurboID to account for the unique subunits of the EARP and GARP, respectively; and VPS52-TurboID and VPS53-TurboID, two shared subunits of the GARP and EARP. All constructs were expressed under the regulation of the COG4 promoter region, allowing expression of tagged CATCHR subunits at near-endogenous levels^22,24^. For each construct, a single clone was selected based on its ability to fully rescue the KO phenotype and to express the TurboID fusion protein at levels comparable to those of the corresponding endogenous protein. One cellular defect associated with depletion of COG complex subunits^26^, as well as depletion of VPS54 and VPS53 (GARP subunits)^10^, is increased cell surface binding of the lectin HPA, which binds to exposed Tn antigen on underglycosylated proteins. As shown in **Figure 1B** and **1C**, this defect is rescued by the TurboID-tagged protein. Depletion of the VPS52 subunit of the GARP and EARP complex, increases levels of the lysosomal protein LAMP2, which is rescued by VPS52-TurboID (**Figure 1B**). Depletion of the EARP subunit VPS50 has been shown to decrease abundance of the other EARP complex subunits^11^. Similarly, VPS50 KO cells exhibit decreased VPS52 and VPS53 abundance, both of which are rescued by VPS50-TurboID (**Figure 1D**). To further validate the functionality of the CATCHR-TurboID constructs, we used immunofluorescence (IF) to assess their localization and labeling region (**Figure 1E**). As expected, both COG subunits and the GARP subunit VPS54-TurboID localize to and biotinylate proteins in the Golgi. Differently, VPS50-TurboID, VPS52-TurboID, and VPS53-TurboID show mostly endosomal localization. This localization is expected for VPS52-TurboID and VPS53-TurboID as these subunits mainly exist as part of the EARP complex^11^. Similar to VPS50, VPS52-TurboID and VPS53-TurboID are recruited to RAB4A-positive endosomes (**Figure 4B** and **Supplementary Figure 1 B-D**). However, closer inspection of VPS52-TurboID and VPS53-TurboID also reveals biotinylated regions in discrete perinuclear puncta that colocalize with the Golgi marker GOLGA4/P230 (**Figure 1E**), which is consistent with these two subunits also being part of the GARP complex. The specificity of proximity labeling by each CATCHR-TurboID construct was determined by comparison with similarly expressed GFP-TurboID, which is diffusely distributed throughout the cytoplasm and does not preferentially localize to any intracellular compartment (**Supplementary Figure 1A**). For this reason, GFP-TurboID works as a background biotinylation control for identifying compartment-specific proximity interactions of peripheral membrane proteins such as the CATCHR complexes. Collectively, these data demonstrate that the CATCHR-TurboID fusion proteins retain biological function, localize correctly within the cell, and enable highly specific mapping of the molecular environments surrounding Golgi and endosomal CATCHR complexes.

### Golgi CATCHRs integrate within functionally analogous trafficking modules

CATCHR-proximal proteins were labeled by short (15 minute) incubation of cells with biotin followed by affinity purification of biotinylated proteins on streptavidin magnetic beads^25^.

Biotinylated proteins were identified by label-free mass spectrometry (MS). The MS proteomics data have been deposited to the ProteomeXchange Consortium via the PRIDE^27^ partner repository with the dataset identifier PXD071676 and 10.6019/PXD071676. The curated datasets are available in **Supplementary Table 1**. Each TurboID proximity-labeling experiment was performed with four to five biological replicates. To assess the specificity and reliability of the MS data, we first examined the enrichment of known subunits of the corresponding tethering complexes. As expected, COG4- and COG6-TurboID preferentially enriched all eight COG subunits, whereas VPS50-, VPS52-, and VPS53-TurboID enriched EARP subunits, and VPS52-, VPS53-, and VPS54-TurboID enriched GARP subunits (**Supplementary Figure 2A**). Interestingly, VPS54-TurboID also enriched several COG subunits, while COG4- and COG6-TurboID enriched the GARP subunits VPS53 and VPS54, suggesting spatial proximity between these Golgi-associated tethering complexes. Furthermore, EARP-associated TurboID constructs selectively enriched VPS33B and VIPAS39, the two subunits of the CHEVI complex, but showed little or no enrichment of components of other endolysosomal tethering complexes, including CORVET, HOPS, and FERARI (**Supplementary Figure 2B**). Together, these results validate the specificity of our proximity-labeling approach and reveal distinct but partially overlapping molecular environments of Golgi and endosomal tethering complexes.

We next assessed whether each CATCHR-TurboID construct preferentially labeled proteins residing in the intracellular compartment where the corresponding complex is known to function. Specifically, we expected the COG and GARP complexes to preferentially label Golgi-associated proteins, whereas EARP was expected to label proteins associated with endosomal membranes. To test this, proteins significantly enriched by each TurboID construct were compared with a curated database of Golgi and endolysosomal proteomes^28,29^. The resulting datasets were visualized as enrichment plots showing the subcellular distribution of the identified proteins (**Figure 2A**, **Supplementary Table 2**). As expected, VPS50-TurboID and VPS52-TurboID showed significant enrichment of endolysosomal proteins, consistent with the established localization of the EARP complex. VPS53-TurboID and VPS54-TurboID labeled both Golgi and endosomal proteins, reflecting the dual role of VPS53 as a shared subunit of GARP and EARP and the dynamic localization of the GARP complex at the TGN-endosome interface. COG4-TurboID significantly enriched Golgi-localized proteins, whereas COG6-TurboID exhibited a similar, although not statistically significant, preference for Golgi proteins. The lack of statistical significance for COG6-TurboID resulted from the strong enrichment of a small number of endolysosomal proteins, including EEA1 and IFT20. Overall, these analyses demonstrate that each TurboID construct selectively labels proteins within the expected intracellular compartment, confirming the high spatial specificity of the proximity-labeling strategy.

**Figure 2.**
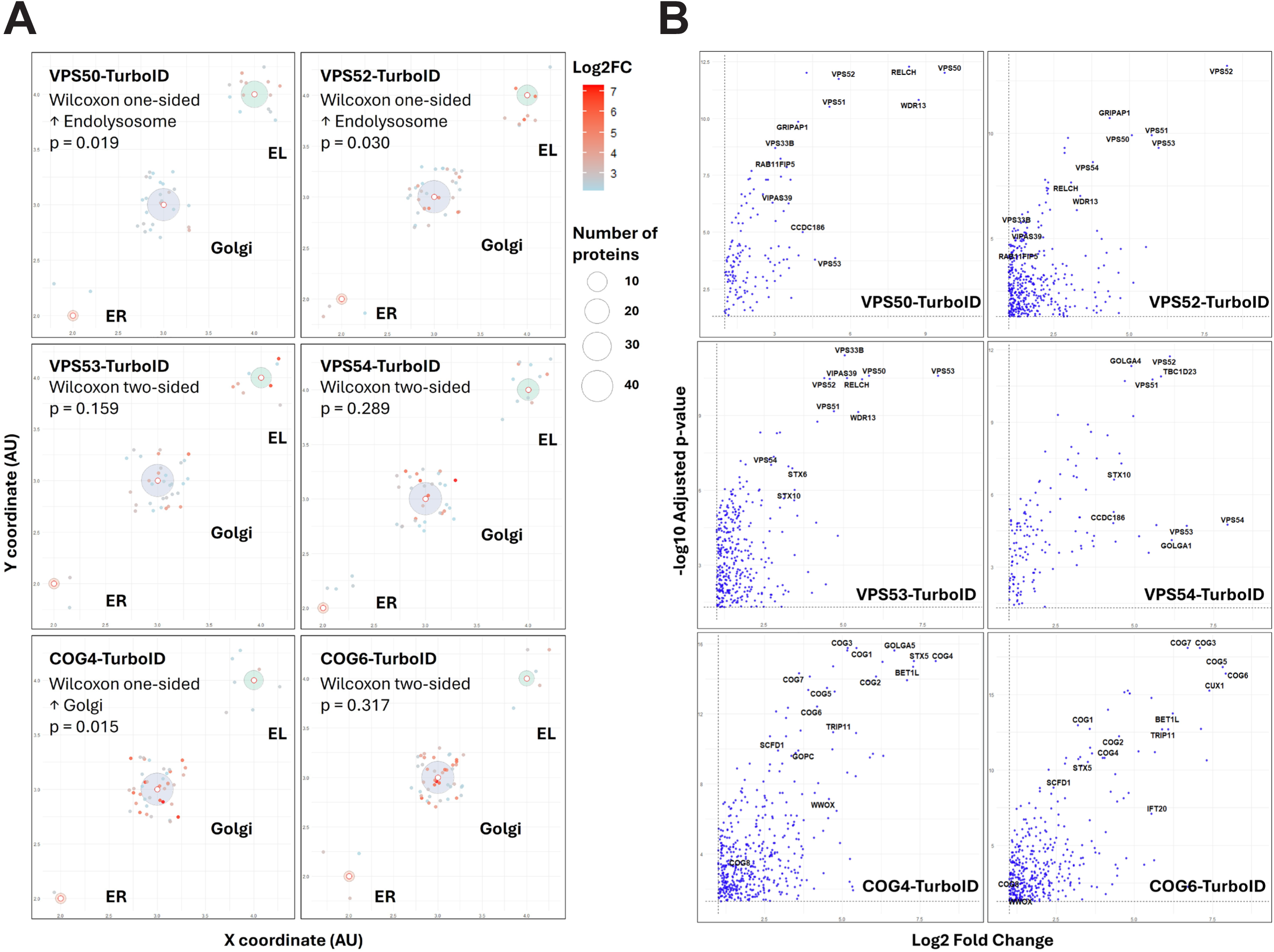
CATCHR-TurboID proximity proteomes show preferential enrichment of compartment-associated proteins and identify established trafficking partners. **(A)** Spatial enrichment analysis of proteins identified across the six CATCHR-TurboID proximity proteomes. Each point represents an individual protein, colored by its log₂ fold enrichment (log₂FC) relative to the soluble GFP-TurboID control. For spatial separation, points are positioned using a k-nearest-neighbor (kNN) algorithm based on their log₂FC profiles. Gray circles indicate protein groups assigned to the Golgi, endolysosomal (EL), or endoplasmic reticulum (ER) compartments, based on curated organellar proteome annotations; circle size reflects the number of proteins assigned to each compartment. VPS50- and VPS52-TurboID show preferential enrichment of endolysosomal proteins, whereas COG4-TurboID shows significant enrichment of Golgi-associated proteins. VPS53- and VPS54-TurboID identify proteins associated with both the Golgi and endolysosomal compartments, consistent with the distribution of GARP and EARP trafficking machinery. No significant enrichment of ER-associated proteins was detected. **(B)** Volcano plots showing protein enrichment in each CATCHR-TurboID proximity proteome relative to the GFP-TurboID control. Each point represents an individual protein. The x-axis indicates log₂ fold enrichment (log₂FC), calculated as the ratio of protein abundance in the indicated CATCHR-TurboID sample to that in the GFP-TurboID control, and the y-axis indicates statistical significance (−log₁₀ p value). Selected significantly enriched proteins are labeled. EARP-associated proximity proteomes identify the complex subunits VPS50, VPS51, VPS52, and VPS53, together with GRIPAP1, CCDC186, VPS33B, VIPAS39, RAB11FIP5, and RELCH. VPS54-TurboID identifies GARP subunits and trafficking-associated proteins, including STX10, TBC1D23, GOLGA1, and CCDC186. COG4- and COG6-TurboID enrich COG subunits, Golgi coiled-coil tethers, SNARE-associated proteins, and WWOX.

Analysis of each of the datasets reveals the enrichment of expected partners as shown in the volcano plots in **Figure 2B**. The highest hits of all subunits of the investigated CATCHRs were components of the trafficking machinery, including CCTs **(Figure 3A),** SNAREs and SM proteins (**Figure 3B**), and other trafficking-related proteins (**Figure 3C**). Western blot (WB) analysis confirmed the specificity of TurboID labeling (**Figure 3D**).

**Figure 3.**
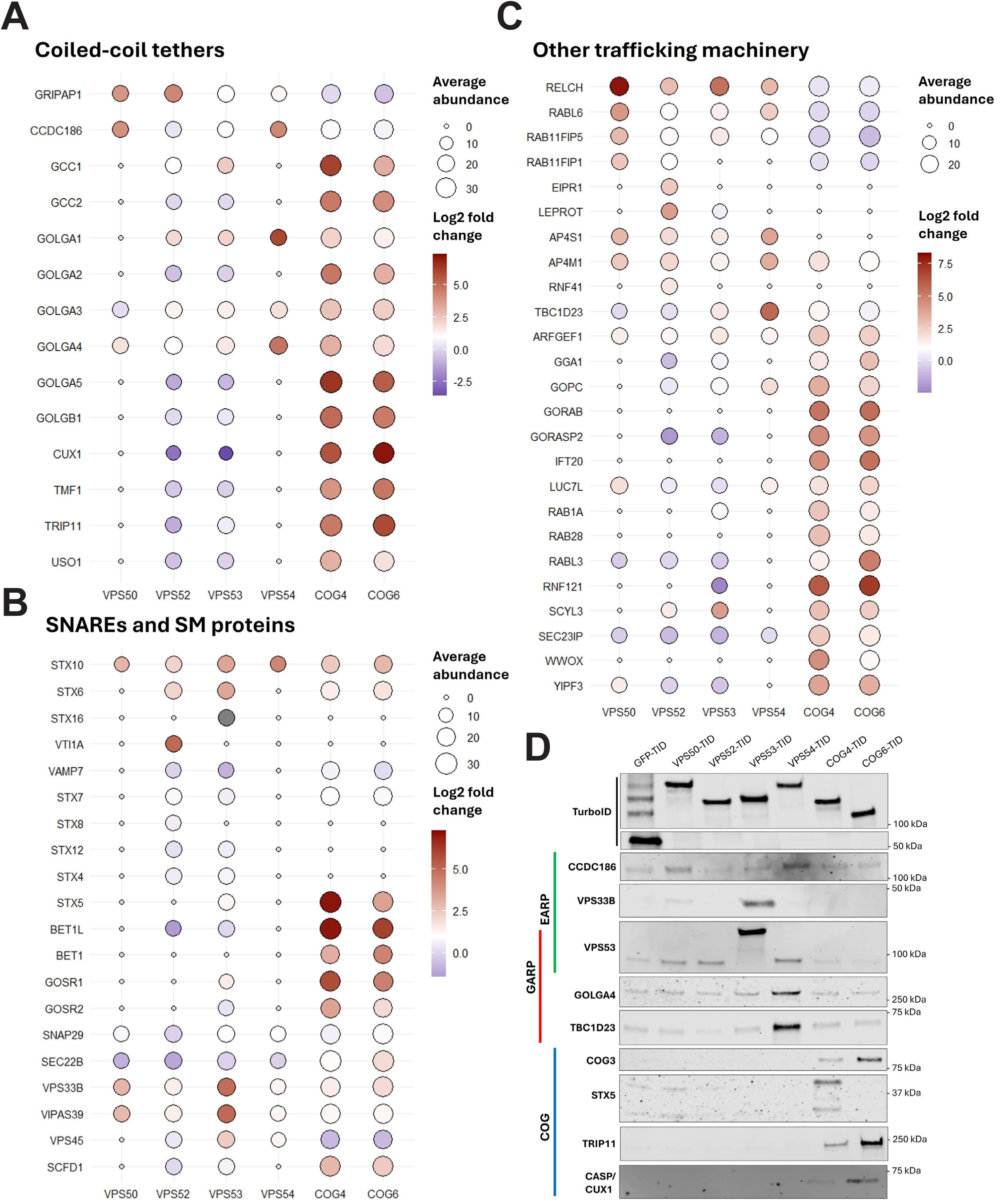
Distinct trafficking modules are associated with Golgi and endolysosomal CATCHR complexes. Dot plots summarize the enrichment of selected trafficking factors identified in the proximity proteomes of VPS50-, VPS52-, VPS53-, VPS54-, COG4-, and COG6-TurboID. Dot size represents average protein abundance, and color indicates log2 fold enrichment relative to GFP-TurboID controls. (**A**) Coiled-coil tethers (CCTs). EARP proximity proteomes are enriched for the endosomal tether GRIPAP1 and the evolutionarily conserved tether CCDC186, whereas GARP preferentially labels CCDC186, GOLGA1, and GOLGA4. COG4- and COG6-TurboID strongly enrich multiple Golgi CCTs, including GOLGA1, GOLGA2/GM130, GOLGA5/golgin-84, GOLGB1/giantin, TMF1, TRIP11/GMAP210, GCC1, GCC2/GCC185, and CUX1/CASP, indicating extensive association of the COG complex with Golgi tethering networks. (**B**) SNAREs and Sec1/Munc18 (SM) proteins. GARP proximity labeling preferentially enriches STX16 and associated SNARE machinery, whereas EARP is enriched for the VPS33B–VIPAS39 CHEVI complex. COG proximity proteomes are enriched for Golgi SNAREs and regulators, including STX5, BET1L, GOSR1, GOSR2, SCFD1, and SNAP29, consistent with a role for COG in regulating intra-Golgi membrane fusion. (**C**) Additional trafficking factors identified in each proximity proteome. EARP preferentially labels RELCH and RAB11FIP5, RAB11-associated machinery involved in recycling endosome function. GARP proximity proteomes are enriched for TBC1D23 and Golgi-associated trafficking factors. COG4- and COG6-TurboID identify multiple regulators of Golgi organization and vesicular transport, including SEC23IP, RNF121, SCYL1, RAB1A, RAB6, and the candidate trafficking regulator WWOX. Together, these data indicate that each CATCHR complex functions within a specialized trafficking module composed of CCTs, SNAREs, SM proteins, and Rab-associated factors. (**D**) Immunoblot validation of selected proximity proteomics hits. Streptavidin pull-downs from TurboID-expressing cells confirm the selective enrichment of CCDC186, VPS33B, VPS53, GOLGA4, TBC1D23, COG3, STX5, TRIP11/GMAP210, and CUX1/CASP by the predicted CATCHR complexes, validating the proteomic datasets.

Previous analysis of COG complex localization and function indicated that COG is associated with rims of multiple Golgi cisternae and orchestrates multiple steps in intra-Golgi trafficking^30,31^. Consistent with its localization, both COG4-TurboID and COG6-TurboID specifically biotinylated all known Golgi-localized CCTs. In addition to previously described COG-interacting CCTs, such as GOLGA5/Golgin-84^32^, CUX1/CASP^13^, GOLGA2/GM130^13^, TMF1^13^, and USO1/p115^33^, our analysis identified several previously unrecognized COG-associated CCTs, including GCC1, GOLGB1/Giantin, TRIP11/GMAP-210, GCC2/GCC185, GOLGA4/Golgin-245, GOLGA3/Golgin-160, and GOLGA1/Golgin-97. Specific proximity labeling of TRIP11 and CASP by both COG4- and COG6-TurboID is shown in **Figure 3D**. These findings substantially expand the known network of potential interactions between the COG complex and CCTs. COG4- and COG6-TurboID labeled Golgi CCTs in a similar, yet distinct, hierarchical pattern, indicating that the two COG lobes occupy overlapping yet spatially distinct molecular environments. In descending order of enrichment, COG4-TurboID preferentially labeled GOLGA5, GCC1, CASP, GOLGB1, GOLGA2, TRIP11, and GCC2, whereas COG6-TurboID showed particularly strong enrichment of CASP, TRIP11, GOLGA5, TMF1, GOLGB1, and GCC2. This graded labeling pattern suggests that COG is embedded within an organized Golgi tethering network, and that COG4- and COG6-containing lobes engage partially distinct subsets of Golgi CCTs.

Similarly, COG-TurboID analysis confirmed the proximity of the COG complex to the Qa-SNARE STX5 and most components of its cognate SNARE machinery, including recently identified partners VAMP7 and SNAP29^34^. In addition to established STX5 partners, COG-TurboID consistently labeled STX6, whereas its canonical partners STX16, VTI1A, and VAMP4 were not detected. This finding is consistent with previously reported co-isolation of STX6 with the COG complex^35^ and raises the possibility that STX6 participates in a previously unrecognized STX5-containing Golgi SNARE complex. Analysis of Golgi SNARE proteins suggested a hierarchy of enrichment within the COG proximity proteomes (**Figure 3B**). STX5 was the highest-ranking SNARE identified by COG4-TurboID, followed by BET1L/GS15, GOSR1/GS28, BET1, and GOSR2/GS27. In contrast, COG6-TurboID preferentially enriched BET1L, GOSR1, and BET1, with STX5 ranking slightly lower but remaining among the most enriched SNAREs. Both proximity proteomes also recovered the cognate SM protein SCFD1 and additional Golgi SNAREs at lower abundance, demonstrating selective enrichment of the STX5-dependent membrane fusion machinery.

In addition to trafficking factors, the COG proximity proteomes identified several proteins involved in Golgi ion homeostasis. Among the most strongly enriched were the Golgi zinc transporters SLC30A5 and SLC30A6^36^, together with the putative Golgi pH regulator GPR89A/B^37^. Proper zinc concentration and luminal pH are essential for Golgi glycosylation and trafficking^38^, raising the possibility that the COG complex contributes to the organization of Golgi ion homeostasis in addition to vesicle tethering and membrane fusion.

Unlike COG, the GARP complex is primarily localized and functions at the TGN^9^. In agreement with GARP localization, members of the TGN trafficking machinery are among the highest hits. It has been shown that the yeast GRIP domain-containing golgin Imh1 interacts with the GARP complex^39^. However, no CCT has been associated with the mammalian GARP to date. We found that the human homologs of Imh1, GOLGA1 and GOLGA4, were among the highest GARP proximity hits, suggesting that the mammalian GARP also interacts with TGN CCTs.

Additionally, GARP-TurboID specifically labeled the TGN SNAREs STX6, STX10 and VTI1A, confirming their functional and physical interaction observed in mammalian^40^ and yeast^14^ cells. The TGN Qa-SNARE STX16 was selectively detected in the GARP-TurboID proximity proteome but was absent in the GFP-TurboID control dataset. This selective labeling supports the specificity of the GARP–STX16 association. However, the absence of STX16 from the control datasets precluded reliable calculation of its relative enrichment and statistical significance using conventional quantitative proteomic analyses. In addition to canonical tethering and fusion factors, the GARP proximity proteomes identified several trafficking regulators that have not previously been directly linked to the GARP complex (**Figure 3C**). Among the highest-ranking proteins were TBC1D23, a TGN adaptor that bridges endosome-derived vesicles to GOLGA1 and GOLGA4 and promotes endosome-to-TGN transport^41^, and CLINT1/EpsinR, a clathrin adaptor involved in AP-1–dependent trafficking between endosomes and TGN^42^. The enrichment of these proteins further defines the molecular environment of the GARP complex and supports its role as a central organizer of TGN trafficking by integrating vesicle tethering with cargo sorting and membrane transport pathways. Specific proximity labeling of GOLGA4 and TBC1D23 by VPS54-TurboID is shown in **Figure 3D**.

To define the trafficking machinery associated with EARP, we next examined the enrichment of CCTs and membrane fusion proteins in the VPS50-, VPS52-, and VPS53-TurboID proximity proteomes (**Figure 3C**). Among proteins involved in membrane tethering, the EARP proximity proteomes preferentially enriched GRIPAP1, CCDC186, and RELCH (**Figure 3A** and **3C**). GRIPAP1 is an established sorting/recycling endosome tether^43^ whereas RELCH has been shown to be involved in recycling endosome tethering to the TGN^44^. Their concurrent enrichment suggests that EARP is embedded within a specialized tethering network operating at the sorting and recycling endosome interface.

Analysis of membrane fusion machinery revealed selective enrichment of the VPS33B-VIPAS39 (CHEVI) complex^45^, identified in all three EARP proximity proteomes (**Figure 3B**). IF analysis confirmed the localization of VPS53 and VPS33B in RAB4-positive endosomal compartments (**Figure 4A-B** and **Supplementary Figure 1E**). Moreover, the endogenous VPS33B was specifically biotinylated by VPS50-TurboID and VPS53-TurboID (**Figure 3B** and **3D**). Since VPS33B is an SM protein that regulates endosomal Qa SNAREs^46^, we investigated whether disrupting the predicted SNARE-binding interface of VPS53 would affect its proximity to VPS33B. To this end, we generated a VPS53 mutant (VPS53mut-TurboID) carrying three substitutions (D473A/K479A/K480A) within its MUN-like domain at a predicted Qa-SNARE-binding interface. We hypothesized that disrupting VPS53–SNARE interactions would alter the proximity of EARP to the SM protein VPS33B. Strikingly, these substitutions abolished VPS33B proximity labeling by VPS53mut-TurboID (**Figure 4C-D**). Importantly, the mutant retained its localization to RAB4-positive endosomes and labeled VPS50 at levels comparable to wild-type VPS53-TurboID, indicating that the mutations did not substantially impair EARP complex assembly or endosomal targeting (**Figure 4C** and **4E**). Together, these findings suggest that the SNARE-interacting region of VPS53 contributes to the functional association of EARP with VPS33B and supports a role for EARP in VPS33B/VIPAS39-associated endosomal trafficking pathways.

**Figure 4.**
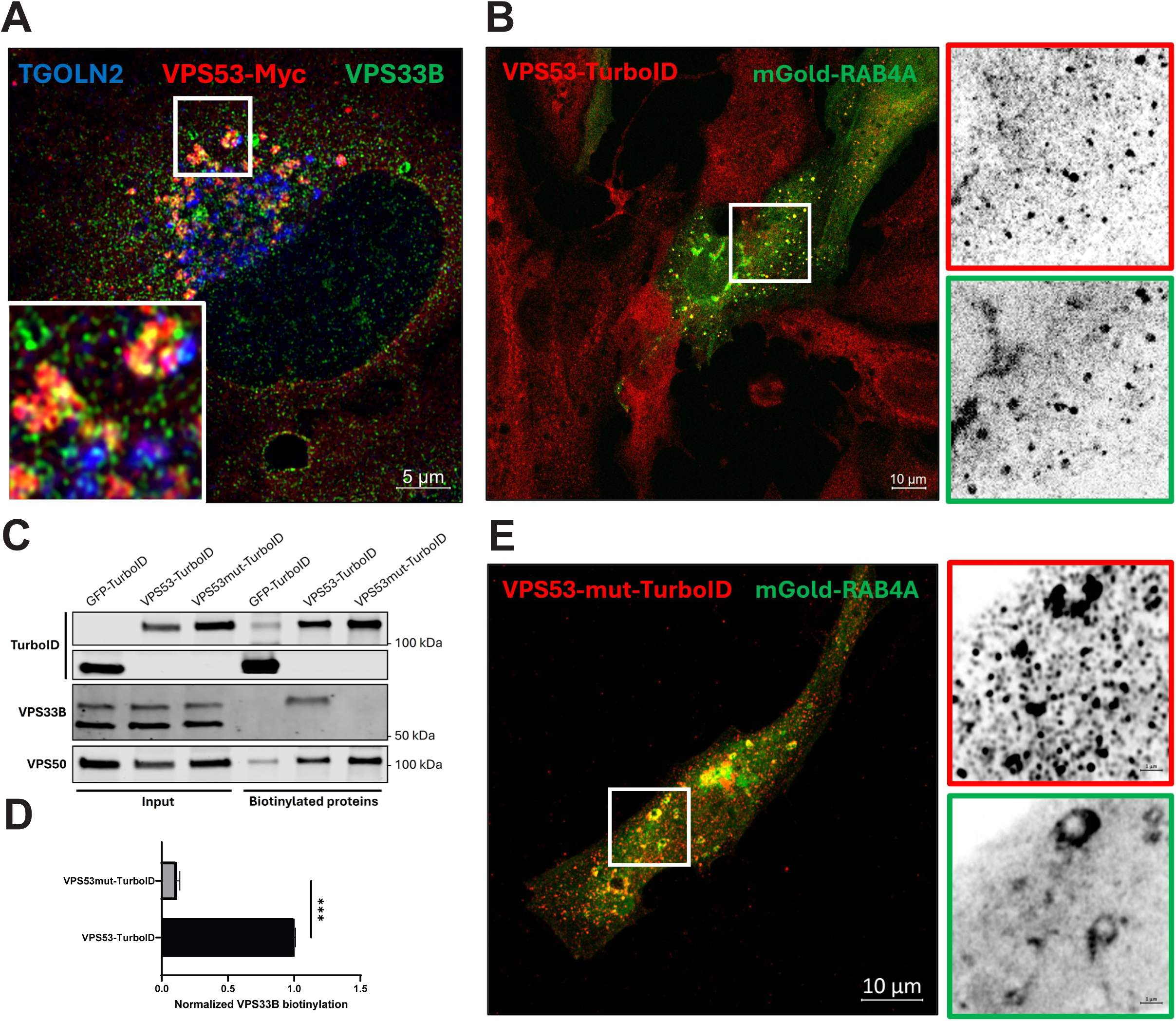
Mutation of the VPS53 MUN-like domain disrupts VPS33B proximity labeling without affecting EARP endosomal localization or assembly. **(A)** IF analysis of myc-tagged VPS53 and endogenous VPS33B in RPE1 cells. VPS53 and VPS33B partially colocalize in peripheral TGN/endosomal structures adjacent to the Golgi. TGOLN2/TGN46 marks the trans-Golgi network (TGN). The boxed region is enlarged in the inset. Scale bar, 5 µm. **(B)** IF analysis of RPE1 cells stably expressing wild-type VPS53-TurboID (red) localizing to mGold-RAB4A-positive endosomes (green). **(C)** Proximity labeling analysis of wild-type VPS53-TurboID and VPS53mut-TurboID. Immunoblotting of input lysates and streptavidin-enriched biotinylated proteins reveals robust labeling of endogenous VPS33B by wild-type VPS53-TurboID, which is markedly reduced by mutations in the VPS53 MUN-like domain. In contrast, VPS50 labeling remains unchanged, consistent with preserved incorporation of mutant VPS53 into the EARP complex. **(D)** Quantification of VPS33B proximity labeling by wild-type and mutant VPS53-TurboID, normalized to wild-type VPS53-TurboID. VPS33B labeling is significantly reduced by mutations in the VPS53 MUN-like domain (\*\**P* < 0.001). **(E)** IF analysis of RPE1 cells stably expressing VPS53mut-TurboID, carrying D473A/K479A/K480A substitutions within the predicted SNARE-binding region of the VPS53 MUN-like domain. VPS53mut-TurboID (red) also localizes to mGold-RAB4A-positive endosomes (green), indicating that the mutations do not disrupt VPS53 endosomal targeting. Boxed regions are enlarged on the right, with individual channels displayed in grayscale. Scale bar, 10 µm. Together, these findings support a functional association between EARP and the CHEVI complex that depends on the predicted SNARE-interacting region of VPS53.

Additionally, EARP preferentially labeled the endosomal Qc-SNARE STX10 alongside multiple components of the endosomal SNARE machinery, consistent with its proposed role in endosomal membrane fusion. Moreover, the EARP proximity proteomes were enriched for proteins associated with cargo sorting, recycling endosome transport, and organelle homeostasis. Among the most prominent were PACS1, a phosphofurin acidic cluster sorting protein that mediates endosome-to-TGN cargo trafficking^47^, and MYO5A, an actin-based motor that functions in Rab11-dependent recycling endosome transport^48^. Interestingly, the VPS52-TurboID screen specifically enriched all V-ATPase subunits, a complex that has been shown to be a TGN pH regulator^49^. Together, these findings further define the molecular environment of the EARP complex and support its role in coordinating cargo sorting, vesicle tethering, organelle homeostasis, and recycling at the sorting/recycling endosome-TGN interface.

Our proximity-labeling analysis also identified several components of Golgi-associated ubiquitination machinery, including previously reported GARP partner E3 ubiquitin ligase RNF41^50^, (**Figure 3C**). Notably, RNF121, another E3 ubiquitin ligase^51^, was among the most strongly enriched proteins in the COG-TurboID datasets (**Figure 3C**). These findings suggest that Golgi-associated CATCHR complexes may functionally interact with components of the ubiquitination machinery. Previous studies have implicated reversible ubiquitination in the regulation of Golgi trafficking proteins, including SNAREs^52,53^. Thus, the selective proximity of RNF41 and RNF121 to GARP and COG, respectively, raises the possibility that ubiquitination contributes to the regulation of CATCHR-dependent membrane trafficking.

Small GTPases and their regulators were also differentially represented across the CATCHR proximity proteomes (**Figure 3C**). The EARP-specific subunit VPS50-TurboID selectively labeled the TGN-localized ARF guanine nucleotide exchange factor ARFGEF1/BIG1, whereas the GARP-specific subunit VPS54-TurboID additionally enriched ARFGEF2/BIG2, RAB1B, and RAB3B, consistent with its broader association with Golgi and TGN trafficking. In contrast, the shared GARP/EARP subunits VPS52- and VPS53-TurboID recovered a more extensive set of GTPase regulators, including ARFGAP1, ARHGEF1, ARFGEF1/2, GBF1, and the small GTPases RAB1A, RAB12, RAB3B, and RAB23, reflecting the common molecular environment shared by the GARP and EARP complexes. The COG proximity proteomes exhibited a distinct profile, with COG4-TurboID preferentially labeling ARFGEF1, ARHGAP10, RAB1A/B, RAB7A, RAB10, RAB11B, RAB28, and Rab-like protein RABL3, whereas COG6-TurboID enriched ARFGAP3, ARFGEF1, ARHGEF2, GBF1, RAB1A, and RABL3. Interestingly, another Rab-like GTPase, RABL6, was detected in both the GARP and EARP proximity proteomes, suggesting that it may shuttle between the TGN and sorting/recycling endosomes, thereby coordinating trafficking at this membrane interface. Together, these findings demonstrate that each CATCHR complex associates with a characteristic repertoire of small GTPases and GTPase regulators, reflecting their distinct functional roles within secretory and endolysosomal trafficking pathways. The lower enrichment of RAB proteins relative to CCTs and SNAREs suggests that Rab GTPases occupy a more peripheral and dynamic position within the CATCHR trafficking modules.

Vesicle coat and cargo-sorting proteins were differentially, but not exclusively, distributed across the CATCHR proximity proteomes. EARP, particularly VPS50-TurboID, showed strong enrichment of AP-3 and AP-4 adaptor subunits, consistent with a recycling endosome-associated sorting environment. GARP/VPS54-TurboID enriched TGN-associated sorting factors, including AP-4 subunits and COPI components. Notably, AP-4 subunits were also recovered in the COG4- and COG6-TurboID datasets, indicating that AP-4 is not EARP/GARP-specific but may represent a broader Golgi/endosomal trafficking module. COG proximity proteomes additionally contained GOPC, SEC23IP/SEC24 proteins, CAV1, and GGA3, suggesting links to COPI/COPII-adjacent and clathrin-associated Golgi trafficking pathways.

Taken together, analysis of the proximity proteome of the CATCHRs suggests that these complexes integrate into higher-order trafficking modules composed of CCTs, SNAREs and SM proteins, as well as trafficking and organellar homeostasis regulators. Defining the molecular modules of the CATCHRs allowed us to mine the datasets for novel members of the trafficking machinery.

### CCDC186 is a vesicular tether working in the GARP domain

CCDC186 was among the most strongly enriched proteins in both GARP- and EARP-TurboID datasets. CCDC186 is an evolutionarily conserved, ubiquitously expressed metazoan protein whose role in membrane trafficking remains incompletely understood. AlphaFold^56^ predictions suggest that CCDC186 forms an elongated, predominantly coiled-coil structure that, when dimerized, resembles known coiled-coil tethers. Given its enrichment in our proximity-labeling datasets and previously reported colocalization with EARP^57,58^, we investigated whether CCDC186 functions as a vesicle tether in GARP- and/or EARP-associated trafficking pathways.

IF analysis revealed that endogenous CCDC186 is predominantly localized to the perinuclear Golgi region, where it partially colocalizes with the GARP-specific subunit VPS54 (**Figure 5A**). To determine its precise position within the Golgi, we treated RPE1 cells with nocodazole to disperse the Golgi ribbon into individual ministacks and examined CCDC186 localization relative to the *cis*-Golgi marker GOLGA2 and the TGN marker TGOLN2/TGN46. CCDC186 was preferentially associated with the TGN region (**Supplementary Figure 3A),** consistent with its proximity to GARP. To test whether CCDC186 can function as a membrane tether, we used a mitochondrial relocalization assay in which CCDC186 was anchored to the outer mitochondrial membrane through either its N- or C-terminus, leaving the opposite terminus exposed to the cytosol. N-terminally anchored CCDC186 (TOM20-CCDC186), which exposes its C-terminal region to the cytosol, induced pronounced mitochondrial clustering (**Supplementary Figure 3B**), a phenotype previously observed with other CCTs in similar assays^59^. TEM further revealed numerous small vesicles, approximately 60 nm in diameter, closely associated with the mitochondrial surface (**Figure 5B**), providing morphological evidence that CCDC186 can recruit and tether vesicular membranes. To investigate the identity of these vesicles, we examined the mitochondrial redistribution of selected endosome-to-TGN trafficking proteins. IF analysis revealed significant recruitment of the cation-independent mannose 6-phosphate receptor (CI-MPR/IGF2R) and the endosomal SNARE STX7 to CCDC186-decorated mitochondria (**Figure 5C-D**). In contrast, redistribution of the cation-dependent mannose 6-phosphate receptor (CD-MPR) and TGOLN2, although an increase was observed, it was not statistically significant (**Figure 5E-F**). The TGN-associated golgin GOLGA4 was also not significantly recruited (**Supplementary Figure 3D**). These findings suggest that CCDC186 preferentially captures a subset of endosome-derived trafficking intermediates rather than indiscriminately recruiting Golgi membranes. Interestingly, reversing the orientation of mitochondrially anchored CCDC186 (CCDC186-OMP), which exposes its N-terminal region to the cytosol, produced a distinct phenotype characterized by smaller mitochondrial aggregates (**Supplementary Figure 3B**). TEM revealed electron-dense connections between adjacent mitochondrial membranes (**Supplementary Figure 3C**), consistent with possible CCDC186-mediated interactions between neighboring mitochondria rather than recruitment of free vesicles. This orientation-dependent behavior suggests that the C-terminal region of CCDC186 is particularly important for vesicle capture.

**Figure 5.**
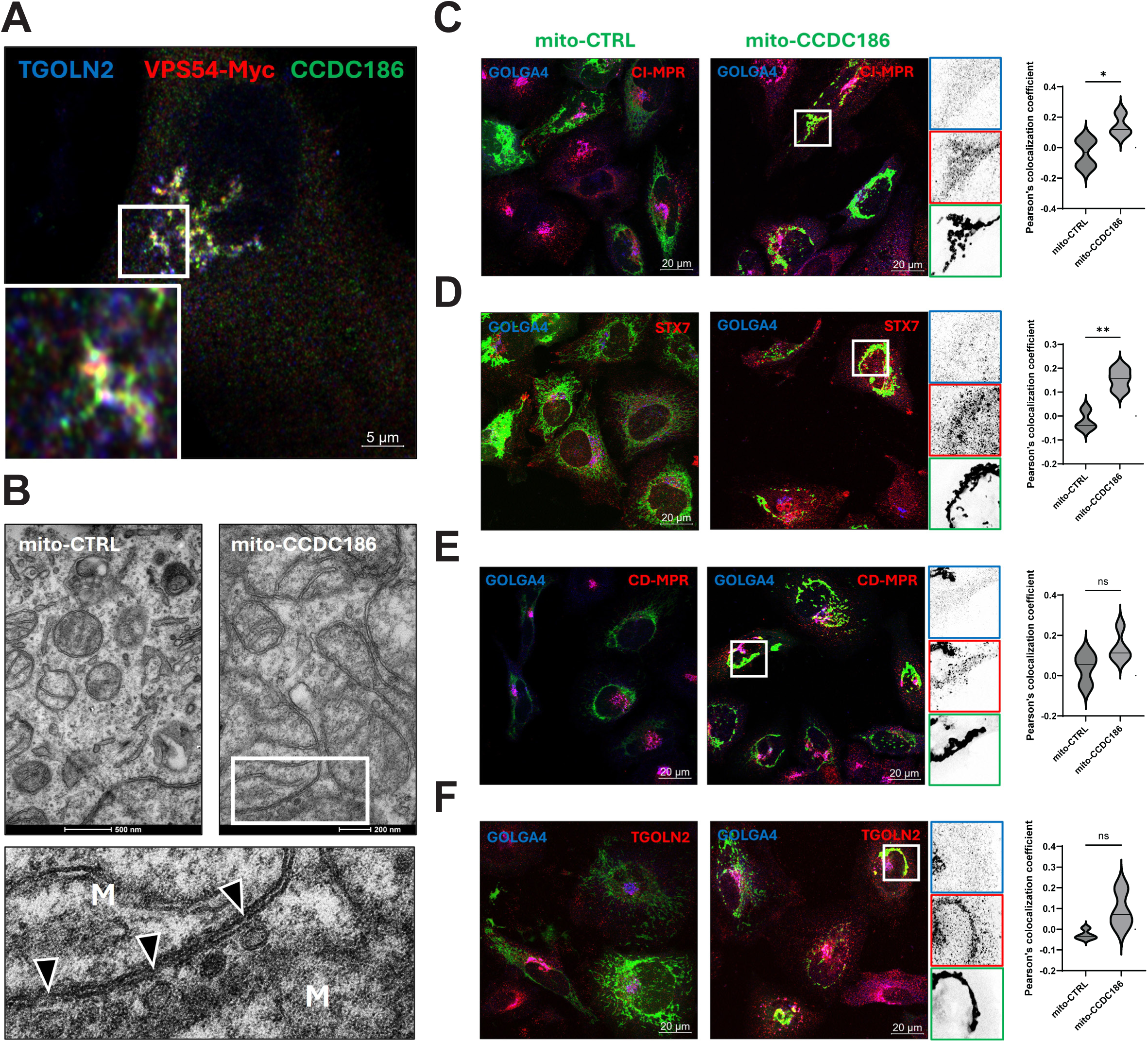
GARP-associated coiled-coil protein CCDC186 tethers endosome-to-TGN transport vesicles. **(A)** IF analysis of myc-tagged VPS54 and endogenous CCDC186 in RPE1 cells. VPS54-positive structures partially colocalize with CCDC186 in TGN-associated compartments, supporting the spatial association of CCDC186 with the GARP complex. TGOLN2/TGN46 marks TGN. The boxed region is enlarged in the inset. Scale bar, 5 µm. **(B)** TEM of cells expressing mitochondrially anchored CCDC186 (mito-CCDC186) or a mitochondrial control construct (mito-CTRL). Mitochondrial recruitment of CCDC186 promotes the accumulation of small vesicles adjacent to mitochondrial membranes, consistent with vesicle tethering activity. The boxed region is enlarged in the lower panel, with arrowheads indicating vesicles associated with mitochondria (M). Scale bars, 200 and 500 nm (upper panels). **(C-F)** IF analysis of intracellular trafficking markers in cells expressing mito-CCDC186 or mito-CTRL (green). Golgi are labeled with GOLGA4 (blue). Recruitment of CCDC186 to mitochondria significantly increases the association of CI-MPR (**C**) and STX7 (**D**) with mitochondrial structures, whereas CD-MPR (**E**) and TGOLN2/TGN46 (**F**) show increased but statistically nonsignificant association. Representative images, enlarged boxed regions, and corresponding Pearson’s colocalization coefficient distributions are shown. Scale bars, 20 µm. Statistical significance is indicated as *P* < 0.05 (*), *P* < 0.01 (**); ns, not significant. Together, these results identify CCDC186 as a vesicle-tethering protein capable of capturing CI-MPR- and STX7-positive transport intermediates, consistent with a role in endosome-to-TGN trafficking.

To further investigate the role of CCDC186 in intracellular trafficking, we depleted endogenous CCDC186 using siRNA and confirmed efficient KD by immunoblotting (**Supplementary Figure 3E-F**). CCDC186 depletion resulted in altered fibronectin secretion, accompanied by intracellular accumulation and increased secretion of pro-cathepsin D (**Supplementary Figure 3G**). Although the pro-cathepsin D secretion phenotype was less pronounced than that observed in VPS54 KO cells, these findings suggest that CCDC186 contributes to endosome-to-TGN trafficking and cargo sorting, potentially functioning alongside the GARP complex.

### WWOX is a novel member of the trafficking machinery working in the COG domain

WWOX (WW domain-containing oxidoreductase) is a multifunctional protein implicated in neuronal development^60,61^ and cancer biology^62^. Evidence shows that downregulation of WWOX is associated with poor prognosis of cancer patients, and different mechanisms have been proposed to explain its pathogenic mode of action^63,64^.

WWOX was consistently enriched in COG-TurboID proximity labeling samples of RPE1 cells (**Figure 2B** and **3C**) and HEK293T cells (data not shown), suggesting a previously unrecognized association with the COG complex. To investigate this interaction, we employed a split-TurboID protein complementation assay^25^. Following transient transfection of RPE1 cells, COG4-splitC-TurboID exhibited a diffuse cellular distribution (**Figure 6A**), potentially reflecting its slow incorporation into the endogenous COG complex, whereas splitN-TurboID-WWOX localized predominantly to the Golgi (**Figure 6B**). Neither construct produced appreciable Golgi-associated biotinylation when expressed individually. In contrast, co-expression of both constructs resulted in robust reconstitution of TurboID activity within the GOLGA2-positive Golgi region (**Figure 6C-D**). Immunoblot analysis confirmed expression of both fusion proteins and increased biotinylation of splitN-TurboID-WWOX upon its co-expression with COG4-splitC-TurboID (**Figure 6E**). Importantly, the L73/G74 split-TurboID system uses low-affinity fragments with minimal spontaneous reassociation^25^, providing strong evidence for a specific molecular association between WWOX and COG4 at the Golgi. As a positive control, co-expression of COG4-splitC-TurboID with splitN-TurboID-STX5, a previously established COG4-interacting SNARE, also produced robust Golgi-associated biotinylation (**Figure 6D-E**). To determine the precise Golgi localization of WWOX, we expressed myc-tagged WWOX because commercially available anti-WWOX antibodies did not produce sufficiently specific IF staining. Myc-WWOX localized to Golgi membranes and exhibited substantial spatial overlap with the COG complex subunit COG8 (**Supplementary Figure 4A**). High-resolution imaging of nocodazole-induced Golgi ministacks further revealed a ring-like distribution of WWOX closely associated with COG8 and the Golgi matrix protein GOLGB1 (**Supplementary Figure 4A-B**). Quantitative analysis of its position along the *cis-trans* Golgi axis placed WWOX predominantly in the medial Golgi region, with a slight bias toward the cis side (localization quotient = 0.27; 20 Golgi regions from three cells; **Figure 6F**). This localization is consistent with the endogenous and endogenously tagged WWOX^65,66^. Together, these observations identify WWOX as a Golgi-associated protein positioned within the COG-associated trafficking environment.

**Figure 6.**
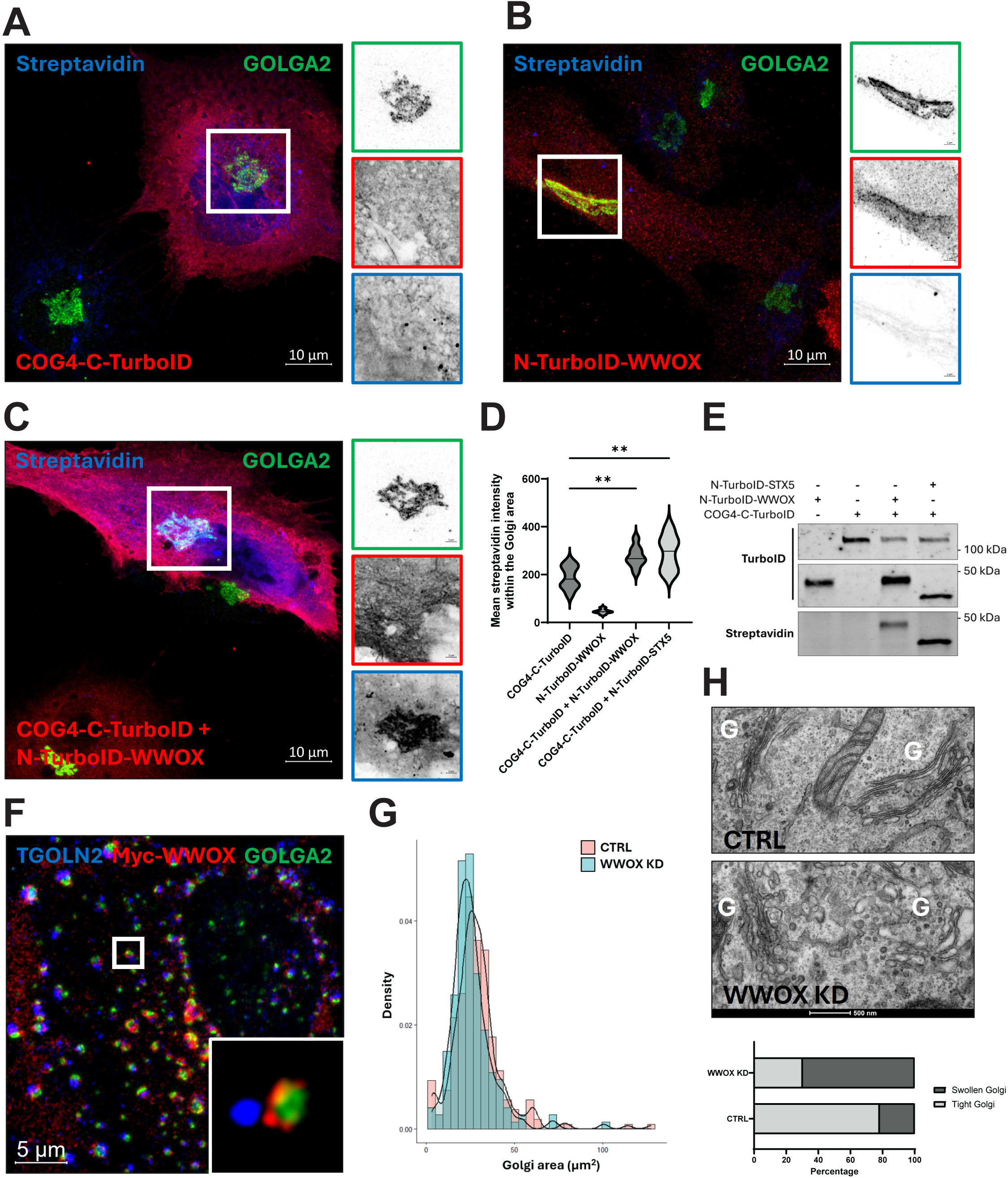
WWOX interacts with the COG complex at the Golgi and is required for normal Golgi organization. (**A-E**) Split-TurboID (red) complementation assay examining the interaction between WWOX and COG4 in RPE1 cells. Cells were transiently transfected with COG4 fused to the C-terminal TurboID fragment (COG4-C-TurboID; **A**), WWOX fused to the N-terminal TurboID fragment (N-TurboID-WWOX; **B**), or both constructs (**C**). At 24 h post-transfection, COG4-C-TurboID exhibited predominantly diffuse cellular localization, whereas N-TurboID-WWOX localized to the Golgi region. Neither construct produced substantial Golgi-associated biotinylation (blue) when expressed individually. In contrast, co-expression of both constructs resulted in robust reconstitution of TurboID activity at GOLGA2/GM130-positive Golgi membranes (green), supporting a specific molecular interaction between WWOX and COG4. Biotinylated proteins were detected using fluorescent streptavidin. Boxed regions are enlarged on the right, with individual channels displayed in grayscale. Scale bars, 10 µm. (**D**) Quantification of streptavidin fluorescence intensity within the Golgi region following expression of individual split-TurboID constructs or their co-expression. Co-expression of COG4-C-TurboID with N-TurboID-WWOX significantly increased Golgi-associated biotinylation, comparable to that observed upon co-expression of COG4-C-TurboID with N-TurboID-STX5, a known COG4-interacting SNARE. Data are presented as violin plots. \*\**P* < 0.01. (**E**) Immunoblot analysis confirming split-TurboID complementation. Cells expressing individual constructs or the indicated combinations were analyzed using anti-TurboID antibodies to detect fusion proteins and streptavidin to detect biotinylated proteins. Increased biotinylation upon co-expression of COG4-C-TurboID with N-TurboID-WWOX or N-TurboID-STX5 confirms functional reconstitution of TurboID activity. (**F**) High-resolution immunofluorescence analysis of myc-WWOX (red) localization relative to the trans-Golgi marker TGOLN2/TGN46 (blue) and the cis-Golgi marker GOLGA2/GM130 (green) at Golgi ministacks in nocodazole-treated RPE1 cells. WWOX localizes predominantly to the medial Golgi, with a localization quotient (LQ) of 0.27 (20 Golgi regions from three cells). The boxed region is enlarged in the inset. Scale bar, 5 µm. (**G**) Quantification of Golgi area in control and WWOX KD RPE1 cells. WWOX depletion significantly reduces Golgi area, indicating altered Golgi organization. Distributions are shown as density plots. Statistical significance was determined using the Wilcoxon rank-sum test with continuity correction (W = 25,487; *P* = 0.00159). (**H**) TEM of control and WWOX-KD cells. WWOX depletion causes pronounced Golgi (G) cisternal swelling and structural abnormalities. Swollen Golgi profiles were observed in approximately 70% of Golgi structures analyzed in WWOX-KD cells (n = 109), compared with approximately 20% in control cells (n = 92). Representative micrographs and quantification of normal versus swollen Golgi profiles are shown. G, Golgi apparatus. Scale bars, 500 nm. Together, these results demonstrate a specific molecular association between WWOX and COG4 at the Golgi and establish an important role for WWOX in maintaining normal Golgi architecture.

To investigate the functional significance of WWOX at the Golgi, we depleted endogenous WWOX using previously validated siRNAs^67^ and confirmed efficient KD by immunoblotting (**Supplementary Figure 4C**). WWOX depletion caused a modest but statistically significant reduction in Golgi area, as assessed by GOLGA2 staining (**Figure 6G** and **Supplementary Figure 4D**). Ultrastructural analysis further revealed pronounced defects in Golgi organization, including swollen cisternae, in approximately 70% of Golgi profiles in WWOX-depleted cells compared with 20% in controls (**Figure 6H**). These findings indicate that WWOX contributes to the maintenance of normal Golgi architecture.

Because disruption of Golgi organization frequently affects glycosylation enzyme localization and function, we next examined whether WWOX depletion altered protein glycosylation. In RPE1 cells, WWOX KD significantly increased cell-surface binding of the lectin HPA, indicating defects in mucin-type O-glycosylation (**Figure 7A-B**). In contrast, binding of the lectin GNL, which recognizes mannose-rich glycans, was not significantly altered in RPE1 cells (**Figure 7B**). Interestingly, WWOX depletion in HEK293 and HeLa cells increased both HPA and GNL binding (**Figure 7C–F**), revealing cell-type-dependent effects on Golgi glycosylation. To confirm that the glycosylation defects were specifically caused by WWOX depletion, we expressed myc-tagged WWOX in WWOX KD HEK293 cells. Importantly, expression of siRNA-resistant myc-WWOX suppressed the increased HPA staining in transfected cells (**Supplementary Figure 4F**), supporting a specific requirement for WWOX in maintaining normal Golgi O-glycosylation. Thus, WWOX is required for normal O-glycosylation across the cell lines examined, whereas its effects on mannose-rich glycan exposure vary between cell types.

**Figure 7.**
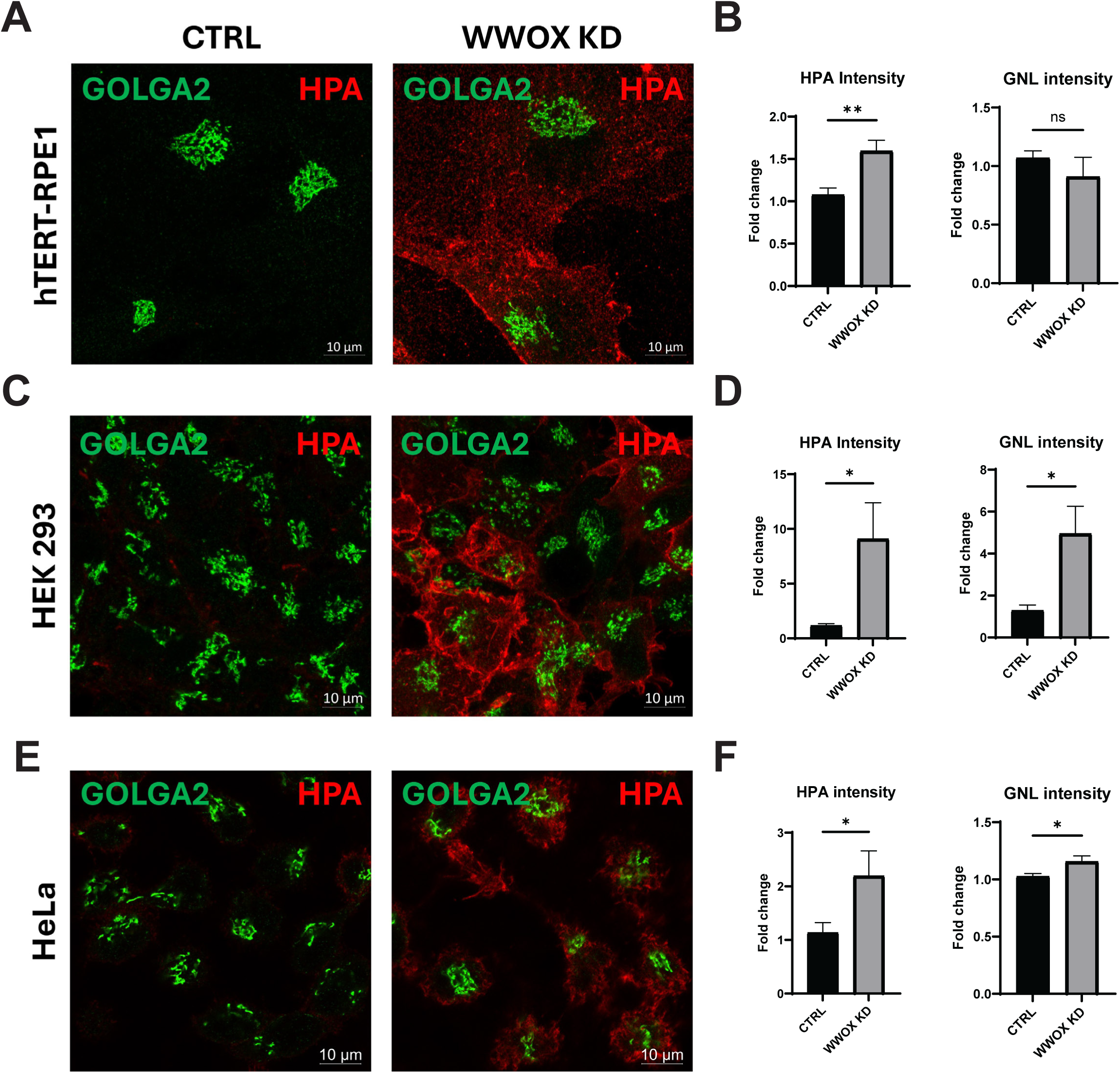
WWOX depletion causes cell-type-dependent defects in N- and O-linked glycosylation. (**A-B**) IF analysis (**A**) and quantification (**B**) of glycosylation defects in hTERT-RPE1 cells following siRNA-mediated WWOX depletion. WWOX KD significantly increases binding of Helix pomatia agglutinin (HPA), which recognizes terminal α-linked N-acetylgalactosamine (Tn antigen), indicating defective O-linked glycosylation. In contrast, binding of Galanthus nivalis lectin (GNL), which recognizes terminal mannose residues, is not significantly altered. (**C-D**) Representative IF images (**C**) and quantification (**D**) of HEK293 cells following WWOX depletion. Both HPA and GNL binding are significantly increased, indicating defects in O- and N-linked glycosylation, respectively. (**E-F**) Representative IF images (**E**) and quantification (**F**) of HeLa cells following WWOX depletion. Similar to HEK293 cells, WWOX KD significantly increases both HPA and GNL binding, consistent with impaired Golgi glycosylation. GOLGA2/GM130 (green) marks the cis-Golgi, and HPA staining is shown in red. Lectin fluorescence intensities are expressed as fold changes relative to control cells. Bar graphs show mean ± SD. Statistical significance is indicated as *P* < 0.05 (*), *P* < 0.01 (**); ns, not significant. Scale bars, 10 µm. Together, these results demonstrate that WWOX is required for normal Golgi glycosylation, with cell-type-dependent effects on N- and O-linked glycan processing.

We also investigated whether WWOX depletion affects the localization of Golgi trafficking machinery. WWOX KD reduced the Golgi association of the COPI coatomer subunit COPB2 relative to the Golgi marker GALNT2 (**Supplementary Figure 4E**). This phenotype is particularly relevant because COPI redistribution has also been observed following acute COG complex inactivation^24^. Together, these findings identify WWOX as a previously unrecognized COG-associated Golgi protein required for normal Golgi architecture, COPI localization, and protein glycosylation. The combination of COG4–WWOX split-TurboID complementation and overlapping loss-of-function phenotypes supports a functional connection between WWOX and COG-dependent membrane trafficking.

### WWOX depletion partially suppresses Golgi protein mislocalization induced by acute COG4 inactivation

The association of WWOX with the COG complex, together with the Golgi defects observed following WWOX depletion, prompted us to investigate whether these proteins functionally interact in maintaining Golgi organization. To address this question, we used RPE1 cells expressing auxin-inducible degron-tagged COG4 (COG4-mAID), allowing rapid depletion of COG4 following treatment with indole-3-acetic acid (IAA)^24,31^. Cells were transfected with control or WWOX-targeting siRNAs for three days, followed by IAA treatment to induce acute COG4 degradation. In control cells, the glycosyltransferase B4GALT1 and the COG complex subunit COG8 were predominantly localized to the Golgi (**Figure 8A**). Acute COG4 depletion caused redistribution of B4GALT1 away from the Golgi and a pronounced loss of Golgi-associated COG8 (**Figure 8B**), consistent with disruption of COG-dependent Golgi trafficking. Interestingly, WWOX KD alone did not substantially alter the Golgi localization of either protein (**Figure 8C**). However, when WWOX-depleted cells were treated with IAA, B4GALT1 localization at the Golgi was partially preserved despite the continued loss of Golgi-associated COG8 (**Figure 8D**).

**Figure 8.**
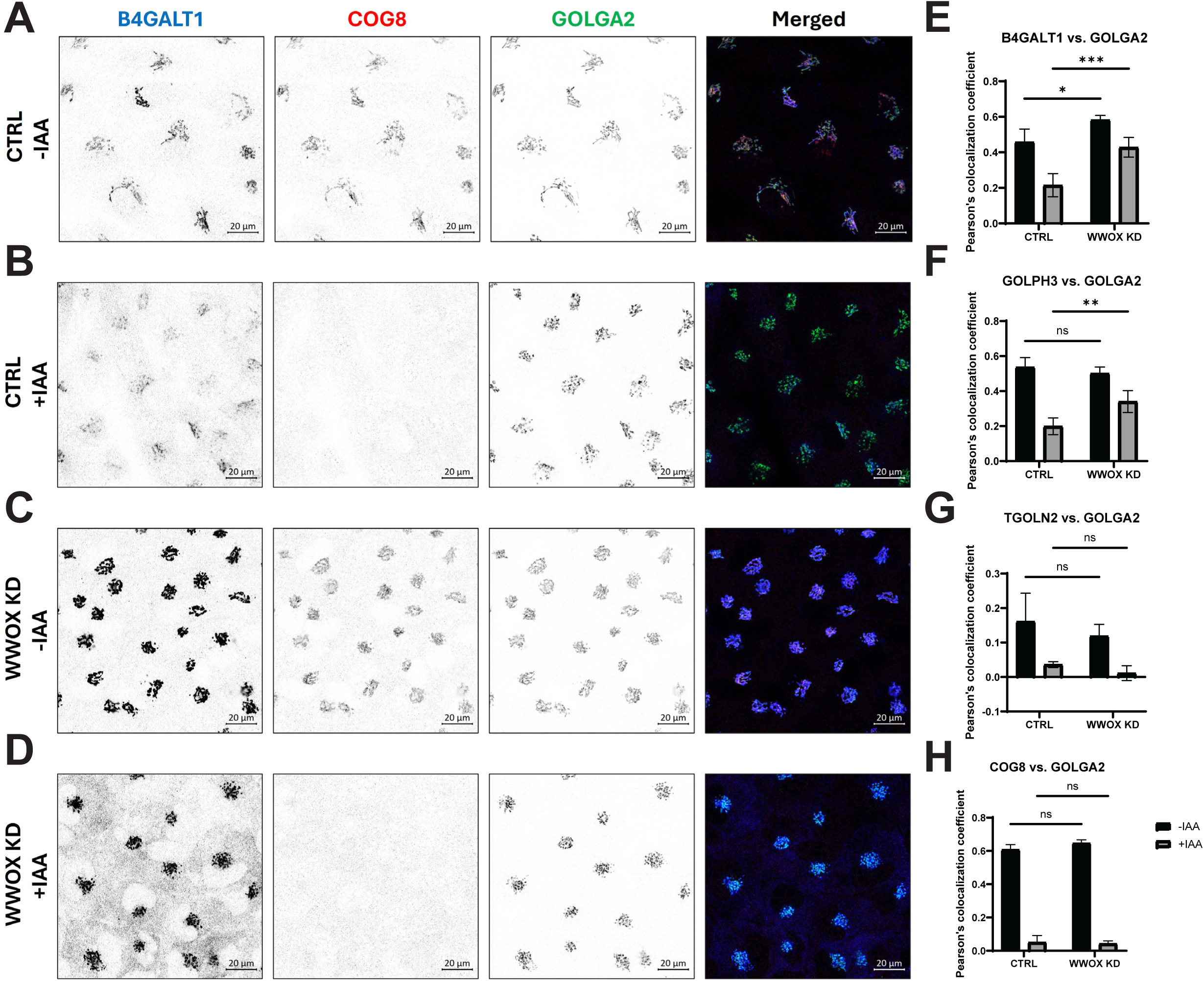
WWOX functionally interacts with the COG complex to regulate Golgi protein localization. (**A-D**) IF analysis of B4GALT1 (blue), COG8 (red), and the cis-Golgi marker GOLGA2/GM130 (green) in hTERT-RPE1 cells expressing COG4 tagged with a C-terminal mini-auxin-inducible degron (COG4-mAID). Cells were treated with control siRNA (CTRL) or WWOX-targeting siRNA (WWOX KD) for three days, followed by treatment with or without 10 µM indole-3-acetic acid (IAA) to induce acute COG4 degradation (∼80% depletion). (**A**) In control cells without IAA, B4GALT1 and COG8 exhibit normal Golgi localization. (**B**) Acute COG4 depletion in control cells causes redistribution of B4GALT1 away from the Golgi and loss of Golgi-associated COG8. (**C**) WWOX depletion alone does not substantially disrupt the Golgi localization of B4GALT1 or COG8. (**D**) Remarkably, combined WWOX KD and acute COG4 depletion preserves substantial B4GALT1 localization at the Golgi, despite the loss of Golgi-associated COG8, indicating that WWOX depletion partially suppresses the B4GALT1 mislocalization phenotype caused by COG4 deficiency. Individual fluorescence channels and merged images are shown. Scale bars, 20 µm. (**E-H**) Quantification of Golgi localization using Pearson’s colocalization coefficients relative to GOLGA2/GM130. (**E**) WWOX depletion significantly attenuates the loss of B4GALT1 from Golgi membranes following acute COG4 depletion. (**F**) A similar partial rescue is observed for the Golgi-associated protein GOLPH3. In contrast, (**G**) TGOLN2/TGN46 and (**H**) COG8 do not exhibit comparable rescue, indicating that the protective effect of WWOX depletion is selective and does not restore COG complex localization. Data are presented as mean ± SD. Statistical significance is indicated as *P* < 0.001 (***), *P* < 0.01 (**), *P* < 0.05 (*); ns, not significant. Together, these results reveal a functional relationship between WWOX and the COG complex in maintaining Golgi protein localization. WWOX depletion selectively attenuates the redistribution of B4GALT1 and GOLPH3 caused by acute COG4 loss without restoring COG8 recruitment to Golgi membranes.

Quantitative colocalization analysis confirmed that WWOX depletion significantly attenuated B4GALT1 redistribution following acute COG4 inactivation (**Figure 8E**). A similar protective effect was observed for GOLPH3, another Golgi-associated protein whose localization is sensitive to COG dysfunction (**Figure 8F**). In contrast, WWOX depletion did not prevent the redistribution of TGOLN2 or the loss of Golgi-associated COG8 following COG4 degradation (**Figure 8G-H**). Thus, the effect of WWOX depletion was selective rather than a general restoration of Golgi organization or COG complex function.

These findings reveal an unexpected functional relationship between WWOX and the COG complex. Although WWOX depletion alone disrupts Golgi morphology and glycosylation, it partially suppresses the mislocalization of selected Golgi proteins caused by acute COG4 inactivation. The persistence of COG8 and TGOLN2 localization defects indicates that this suppression does not reflect a complete rescue of COG-dependent trafficking. Instead, our results suggest that WWOX influences a subset of Golgi trafficking or retention pathways that become particularly important when COG function is compromised

## Discussion

Our work provides the first comprehensive characterization of the *in vivo* molecular landscape of three Golgi-endosomal CATCHR complexes: the COG, GARP, and EARP. Using cell lines stably expressing CATCHR-TurboID constructs at near-endogenous levels, we define the molecular environment of these evolutionarily conserved complexes. The specificity of the proximity proteomic datasets was supported by the correct localization of the TurboID-tagged CATCHR subunits, preservation of their biological function as demonstrated by rescue of the corresponding KO phenotypes, and short labeling times that minimized secondary and nonspecific biotinylation. Together, these features enabled the identification of proteins residing in the immediate molecular environment of each CATCHR complex.

As expected, the most strongly enriched proteins included validated intra- and inter-complex partners. Each TurboID-tagged subunit specifically biotinylated the other subunits of its corresponding complex (**Supplementary Figure 2A**). Remarkably, proximity labeling occurred preferentially within individual COG lobes rather than across the entire COG complex. COG1, COG2, and COG3 were among the most strongly enriched proteins in the COG4-TurboID proximity proteome, whereas COG5 and COG7 were the dominant hits in the COG6-TurboID dataset. Given that all COG subunits are components of a single octameric tethering complex, this differential labeling underscores the spatial discrimination of the TurboID approach.

Moreover, these findings provide *in vivo* support for our model that the COG complex is organized into two subcomplexes (lobes A and B) whose interactions are regulated rather than constitutively fixed^68^.

GARP-TurboID also labeled STX16, GOLGA1, and RNF41, previously characterized GARP-associated proteins in yeast or humans^14,39,50^. The enrichment of these established partners provided an important internal benchmark supporting the sensitivity and specificity of our proximity-labeling strategy.

Intriguingly, reciprocal proximity labeling between COG and GARP components was observed, with COG-TurboID labeling VPS53 and VPS54 and VPS54-TurboID labeling multiple COG subunits (**Supplementary Figure 2A**). These findings raise the possibility that Golgi CATCHR complexes do not function as entirely independent entities. One possibility is that COG and GARP cooperate to regulate STX16-dependent membrane fusion events^69^. Alternatively, the two complexes may form hybrid CATCHR assemblies, expanding the repertoire of tethering and fusion machinery available at the Golgi. Such mixed assemblies could account for the unexpected redundancy of COG lobe B subunits reported in yeast^12,70^ and Drosophila^71,72^.

Interestingly, a hybrid assembly involving RINT1, a subunit of the NRZ complex, and COG has previously been proposed^73^. Consistent with this possibility, RINT1 was detected in the COG4-TurboID proximity proteome, although its enrichment was not statistically significant compared with the GFP-TurboID control. Thus, while our findings raise the possibility of functional cooperation or hybrid assembly among CATCHR complexes, direct biochemical evidence is needed to distinguish hybrid complex formation from the close spatial association of independently assembled complexes.

One of the most striking features of the CATCHR proximity proteomes was the hierarchical organization of associated trafficking factors. Comparison of the proximity proteomes revealed that each CATCHR complex is associated with a distinct but evolutionarily conserved trafficking module composed of CCTs, SNAREs and SM proteins, Rab GTPases and their effectors, vesicular coats, and regulators of trafficking and organellar homeostasis. Among proteins external to the COG complex, Golgi CCTs constituted the most strongly enriched group, with previously established COG interactors, including GOLGA5, GOLGA2, GOLGB1, and TMF1, ranking among the highest hits, whereas TRIP11, GCC1, GCC2, and CASP showed progressively lower enrichment. Components of the STX5-dependent membrane fusion machinery formed the next tier of the proximity proteome. Notably, STX5 was the highest-ranking SNARE identified by COG4-TurboID, whereas COG6-TurboID preferentially enriched vesicular components of the cognate SNARE complex, including BET1L, GOSR1, BET1, and GOSR2. Together with previous biochemical studies demonstrating direct interaction between COG4 and STX5^15,34,74–76^, these findings suggest that the two COG lobes occupy distinct but complementary positions within the membrane fusion machinery.

Alongside several established TGN CCTs, the GARP proximity proteome identified the coiled-coil protein CCDC186, for which we provide direct evidence of vesicle-tethering activity in mammalian cells. The *C. elegans* homolog of CCDC186, CCCP-1, colocalizes with the EARP complex and is required for dense-core vesicle^77^ biogenesis, whereas the *D. melanogaster* homolog, Golgin104, functions as a Rab2 effector associated with the Golgi^78^. In addition, the EARP proximity proteomes recovered EIPR1/TSSC1, a protein previously shown to function together with CCDC186 in dense-core vesicle cargo sorting^58^. The enrichment of CCDC186 and EIPR1 supports the biological relevance of our proximity proteomic datasets and identifies a conserved trafficking module operating at the TGN-endosome interface. Together, these observations indicate that CCDC186 is an evolutionarily conserved CCT that functions predominantly at the TGN while dynamically associating with endosomal membranes, where it cooperates with the EARP complex. Future studies should define the molecular mechanisms governing the dynamic localization of CCDC186 and determine how this CCT, together with EIPR1, coordinates GARP- and EARP-dependent vesicle tethering and cargo transport at the TGN-endosome interface.

Our mitochondrial relocalization experiments provide additional functional evidence that CCDC186 can capture transport intermediates. Mitochondrially anchored CCDC186 induced the accumulation of small vesicles near mitochondria and selectively recruited CI-MPR- and STX7-positive membranes, consistent with the capture of endosome-derived transport intermediates (**Figure 5**). These findings support a role for CCDC186 in endosome-to-TGN trafficking and suggest that its C-terminal region contributes to vesicle recognition or capture. Together, our observations identify CCDC186 as an evolutionarily conserved CCT that functions predominantly at the TGN while potentially associating with endosomal trafficking machinery. Future studies should define the molecular mechanisms governing CCDC186 localization and vesicle recognition and determine how CCDC186 and EIPR1 coordinate GARP- and EARP-dependent transport at the TGN–endosome interface.

Another important finding was the identification of the SM protein VPS33B and its partner VIPAS39, which together form the CHEVI complex, among the most strongly enriched proteins in the EARP proximity proteome. Our results suggest that EARP may function within a larger trafficking module containing CHEVI. Importantly, mutations in the MUN-like domain of VPS53 markedly reduced VPS33B proximity labeling without disrupting EARP assembly or its localization to RAB4-positive endosomes (**Figure 4**). This separation of EARP localization and VPS33B proximity suggests that the VPS53 MUN-like domain contributes to the organization of the EARP-associated fusion machinery. Moreover, RAB11FIP5, a Rab11 effector involved in recycling endosome trafficking and a component of the FERARI complex^79^, was also strongly enriched in the EARP proximity proteome. Together, these observations support a model in which EARP cooperates with CHEVI and Rab11-associated factors to coordinate tethering and fusion events at recycling endosomes, forming a trafficking module different than other endolysosomal tethering complexes (**Supplementary Figure 2B**). Future studies should determine whether EARP and CHEVI interact directly and how their activities are coordinated during endosomal membrane fusion.

An unexpected finding was the enrichment of multiple vesicle coat proteins and coat-associated adaptors within the CATCHR proximity proteomes. Although the specific repertoire differed among COG, GARP, and EARP, components of the COPI, AP-1, AP-3, AP-4, and GGA coat machinery were detected in close proximity to one or more CATCHR complexes. These observations suggest two models that are not mutually exclusive. First, remnants of vesicle coats may remain associated with trafficking intermediates after budding, allowing CATCHR complexes to recognize coat proteins as an additional determinant of vesicle identity during tethering. Such a mechanism would complement the established roles of Rab GTPases, CCTs, and SNARE proteins in specifying vesicle targeting. Alternatively, vesicle docking and fusion sites may be positioned immediately adjacent to vesicle budding domains within Golgi and endosomal membranes. In this model, CATCHR complexes would not only mediate vesicle tethering but also spatially coordinate vesicle formation, docking, and fusion, thereby coupling successive stages of membrane trafficking. Future studies combining high-resolution imaging and functional perturbation of coat components will be required to distinguish between these models.

Importantly, the proximity proteomes identified several regulatory proteins not previously linked to the CATCHRs. These include the transmembrane RING finger E3 ubiquitin ligase RNF121^51,80^, the oxidoreductase WWOX, the Golgi zinc transporters SLC30A5 and SLC30A6, and the putative Golgi pH regulator GPR89, all detected in the COG-TurboID screen. The identification of RNF121 and WWOX raises the possibility that the COG complex associates with regulatory enzymes in addition to canonical tethering and fusion factors, thereby linking membrane trafficking to post-translational regulatory and signaling pathways. Likewise, the enrichment of SLC30A5/6 and GPR89 suggests a potential relationship between the COG complex and the physicochemical environment of the Golgi, where ion homeostasis and luminal pH are essential for glycosyltransferase activity. It is also possible that the CATCHR complexes recognize ion transporters and pH sensors on acceptor membranes as additional determinants of compartment identity. Similarly, VPS52, a shared subunit of the GARP and EARP, may participate in such recognition, as VPS52-TurboID detected RNF41 and all subunits of the V-ATPase complex. Together, these findings indicate that individual CATCHR complexes associate with distinct accessory factors that tailor their activity to specific trafficking pathways. Future studies should determine how these regulatory proteins cooperate with CATCHR complexes and whether disruption of these networks contributes to the glycosylation and trafficking defects characteristic of COG-, GARP-, and EARP-associated disorders.

Among the newly identified COG-associated proteins, WWOX emerged as a particularly compelling candidate regulator of Golgi function. WWOX localized predominantly to the medial Golgi, where it partially colocalized with the COG complex. Importantly, complementation of the low-affinity L73/G74 split-TurboID fragments fused to WWOX and COG4 produced robust Golgi-associated biotinylation, providing strong evidence for a specific molecular association between these proteins (**Figure 6**). This finding independently supports the identification of WWOX in the COG proximity proteome and suggests that WWOX is positioned within the COG-associated trafficking machinery. WWOX depletion caused Golgi fragmentation, defects in O-glycosylation, and partial mislocalization of the COPI coat; phenotypes that closely resemble those observed following COG depletion (**Figures 7** and **8**). The glycosylation phenotypes were not identical across the cell lines examined: WWOX depletion increased HPA binding in RPE1, HEK293, and HeLa cells, whereas increased GNL binding was detected in HEK293 and HeLa cells. These differences suggest that the contribution of WWOX to Golgi glycosylation may depend on cell-specific trafficking requirements or the relative sensitivity of different glycosylation pathways. Consistent with a role in Golgi trafficking, WWOX was previously shown to interact with the ER-Golgi trafficking factor SEC23IP and the Golgi CCT TMF1^81^, both of which were also identified in our COG proximity proteome (**Figure 3A** and **3C**). Unexpectedly, WWOX depletion partially suppressed the mislocalization of selected Golgi proteins following acute COG4 inactivation (**Figure 8**). Specifically, WWOX KD partially preserved the Golgi localization of B4GALT1 and GOLPH3 despite COG4 degradation, but did not prevent TGOLN2 redistribution or the loss of Golgi-associated COG8. This selective suppression suggests that WWOX regulates trafficking pathways that functionally intersect with COG-dependent membrane recycling. One possibility is that WWOX promotes the incorporation of a subset of Golgi-resident proteins into recycling intermediates. Its depletion could therefore reduce their redistribution when COG-dependent vesicle tethering and fusion are impaired.

Alternatively, WWOX may influence COPI-dependent trafficking or other processes involved in maintaining Golgi organization. Importantly, preservation of Golgi localization does not necessarily indicate restoration of normal trafficking or glycosylation. Thus, the functional relationship between WWOX and COG appears more complex than a simple positive regulatory interaction.

Although the molecular mechanism of WWOX remains unknown^60,82,83^, its tandem WW domains could function as a scaffold to recruit PPxY-containing trafficking proteins to Golgi membranes^66,82^, whereas its C-terminal short-chain dehydrogenase/reductase (SDR) domain may regulate Golgi physiology through local NAD(H)- or NADP(H)-dependent redox reactions^84^.

Future studies should define the molecular basis of the WWOX–COG association and determine how WWOX influences Golgi membrane recycling and glycosylation. More broadly, our findings establish an unexpected connection between WWOX and the Golgi trafficking machinery, providing a potential link between defects in membrane trafficking and the neurodevelopmental disorders and cancers associated with WWOX dysfunction. Although TurboID labeling is influenced by protein abundance, lysine accessibility, and interaction dynamics, the reproducible hierarchy observed across all three CATCHR complexes suggests that it reflects the spatial organization of specialized trafficking modules. Remarkably, this hierarchy is consistent with the proposed sequence of events during vesicle tethering, beginning with Rab-mediated vesicle targeting, followed by long-range capture by CCTs and culminating in CATCHR-dependent assembly of the cognate SNARE fusion machinery. Together, these findings support a model (**Figure 9**) in which CATCHR complexes function as central organizing hubs that coordinate successive stages of vesicle tethering and membrane fusion while establishing compartment-specific identity through selective association with CCTs, SNAREs, SM proteins, and pathway-specific regulatory factors. Collectively, our work provides a comprehensive molecular framework for the organization of Golgi and endosomal CATCHR complexes and reveals a fundamental principle of membrane trafficking: rather than functioning as isolated vesicle tethers, CATCHR complexes assemble specialized trafficking modules that integrate long-range capture by CCTs, SNARE-dependent membrane fusion, and acceptor membrane molecular cues for selective fusion. By defining these molecular networks and identifying previously unrecognized regulatory components, our study provides a foundation for elucidating the mechanisms that maintain Golgi and endosomal homeostasis and how their disruption contributes to congenital disorders of glycosylation, neurodevelopmental diseases, and cancer.

**Figure 9.**
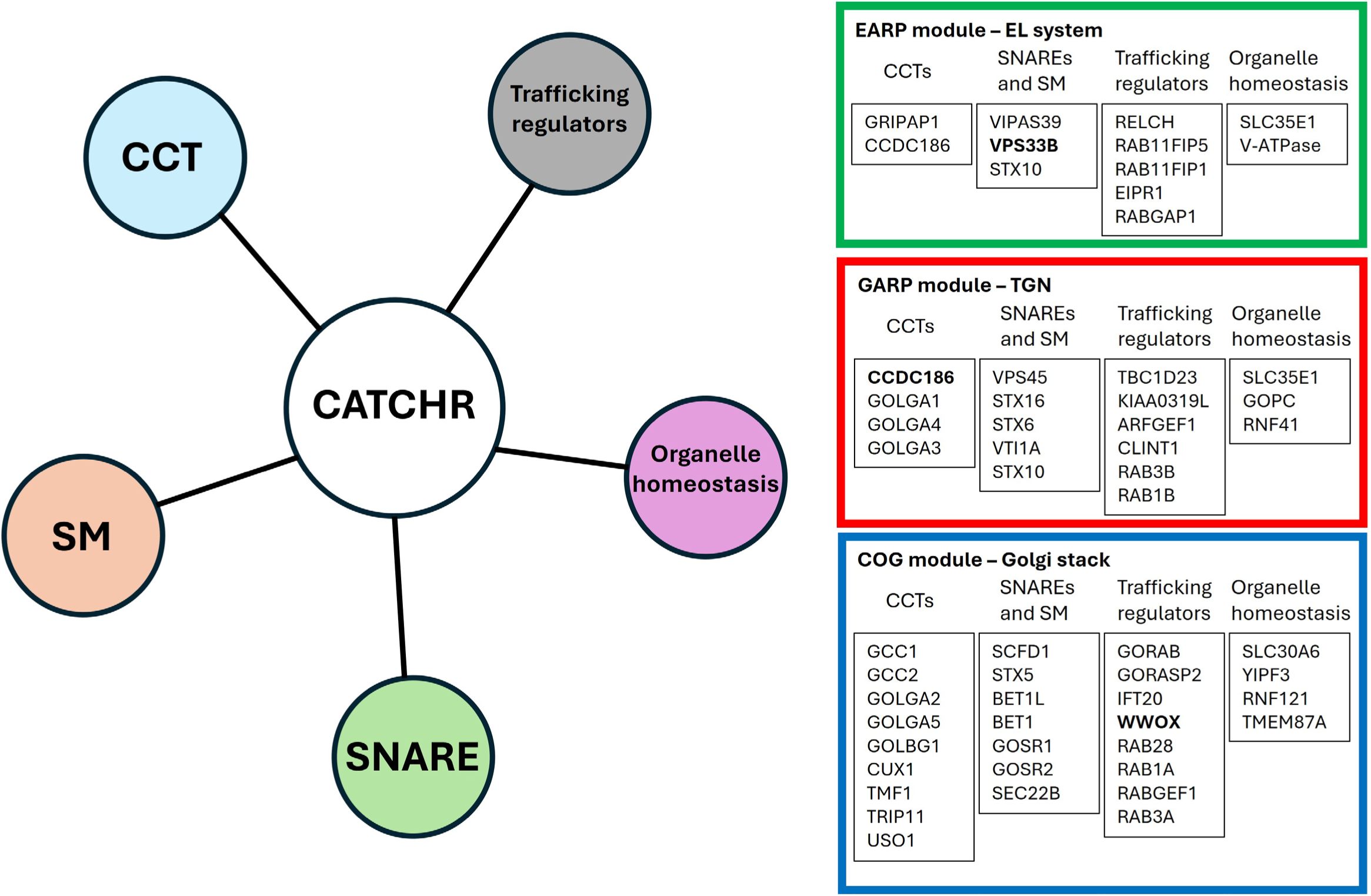
Golgi–endosomal CATCHR complexes function as organizing hubs for vesicle tethering and fusion. Schematic model illustrating the distinct but overlapping trafficking modules organized by the CATCHR complexes EARP, GARP, and COG. Each complex associates with a characteristic network of coiled-coil tethers (CCTs), SNAREs and Sec1/Munc18 (SM) proteins, trafficking regulators, and organelle homeostasis factors. **EARP** (green) operates primarily in the endosomal recycling system and is associated with the VPS33B–VIPAS39 (CHEVI) complex, the SNARE STX10, CCTs GRIPAP1 and CCDC186, and trafficking regulators including RELCH and RAB11FIP5. **GARP** (red) functions at the trans-Golgi network (TGN) and is associated with CCDC186, Golgi-associated CCTs, the STX16–VPS45 fusion machinery, and trafficking regulators including TBC1D23. **COG** (blue) operates primarily at the Golgi stack and is associated with multiple golgins, STX5–SCFD1-dependent fusion machinery, Rab GTPases and their effectors, and the newly identified Golgi-associated regulator WWOX. The proteins shown represent selected components identified through proximity proteomics and functional analyses. Together, these findings support a model in which CATCHR complexes coordinate specialized molecular networks that link vesicle recognition and tethering to SNARE-mediated membrane fusion and organelle homeostasis.

## Materials and methods

### Cell line creation and cell culture

Retinal Pigment Epithelial (hTERT-RPE1), HeLa WT, and HEK293T WT cells used for all experiments were purchased from ATCC. WT and KO cell lines were maintained in DMEM/F12 supplemented with 10% FBS. TurboID-expressing cells were maintained in biotin-free DMEM supplemented with 10% FBS and 10 μg/ml of avidin. Cells were incubated in a 37°C incubator with 5% CO2 and 90% humidity.

RPE1 KO cell lines were created as described in previously^85–87^. These cells were rescued using lentiviral transduction of CATCHR-TurboID constructs. Lentiviruses were made in-house as described previously^85,86^. The plasmids used for lentivirus formation along with the packaged plasmids containing the CATCHR-TurboID sequences are listed below (**Table 1)**. All constructs were cloned into a destination vector containing a G418 resistance gene and the COG4 promoter previously described^88^. After selection with 500 μg/ml of G418, the cells were single-cell sorted, and individual clones were characterized by WB and selected based on the expression level of CATCHR-TurboID hybrid closest to the levels of the endogenous protein.

**Table 1.**
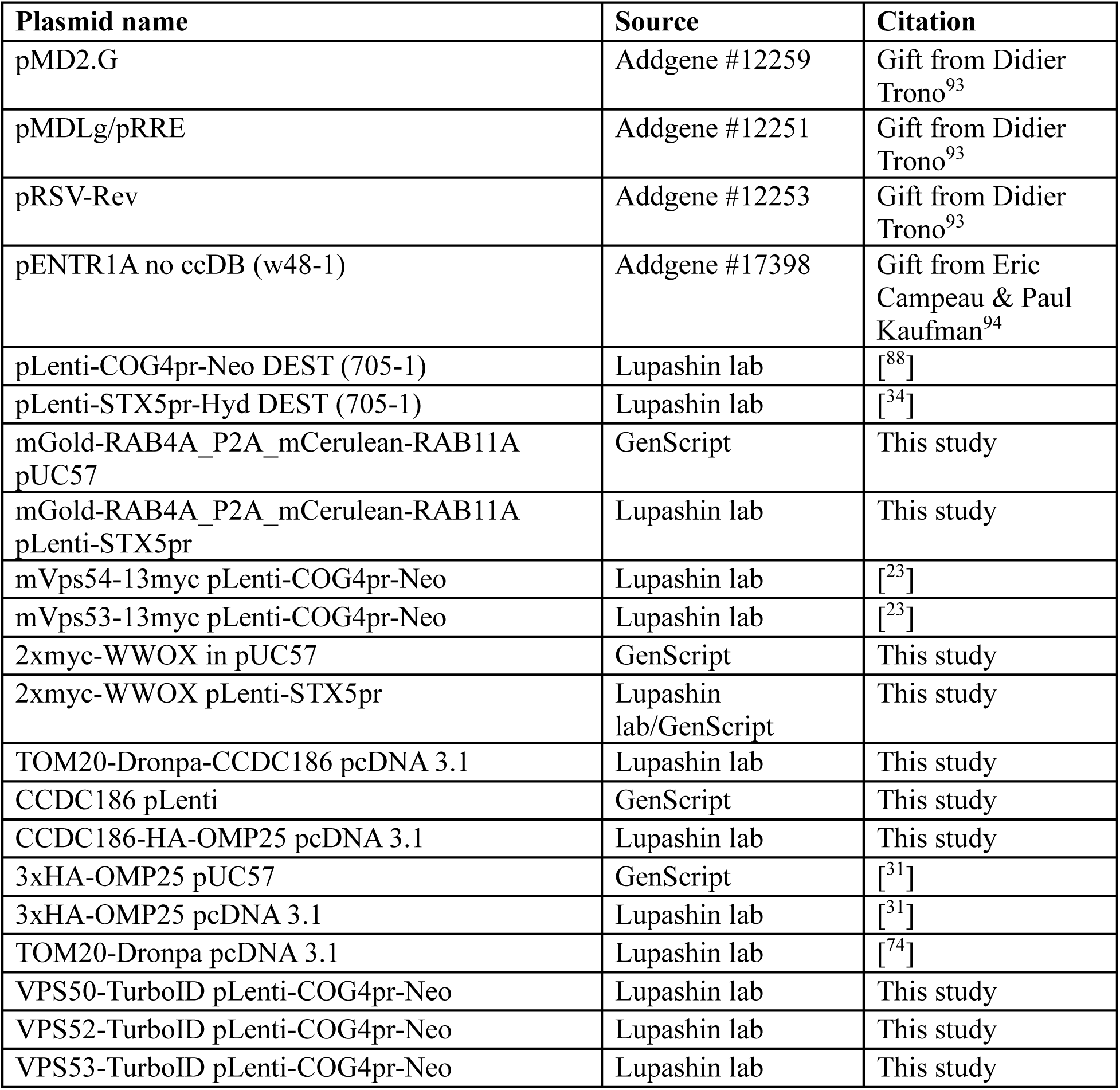
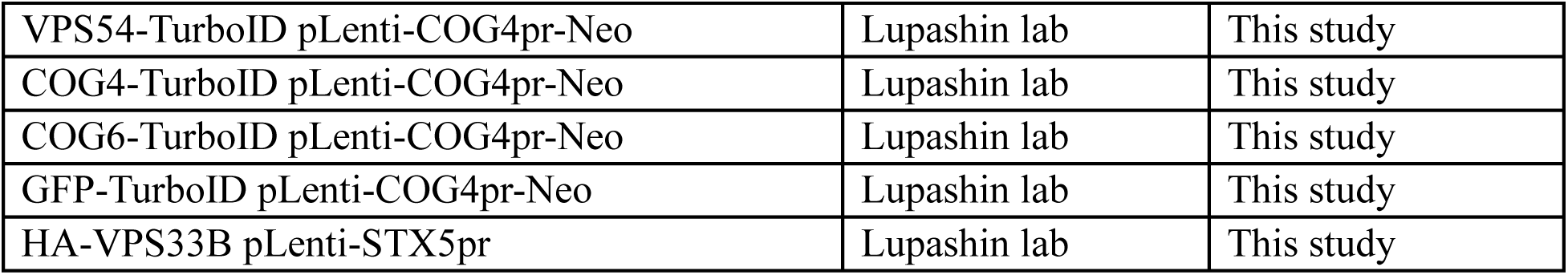
DNA constructs.

### Proximity labeling assays

Cells grown in a 15 cm dish (MS) or 10 cm dish (WB) maintained in biotin-free media for at least 72 h. To initiate biotinylation, cells were incubated with media supplemented with 100 μM biotin for 15 min for MS analysis, 30 min for WB and IF.

For MS, after biotinylation cells were placed on ice and washed four times with ice-cold PBS. Cells were then lysed with ice-cold RIPA buffer supplemented with 100 μM PMSF and Halt protease inhibitor as per the manufacturer’s instructions. Lysate was centrifuged at 4°C for 10 min at 13,000 g to pellet cell debris, the resulting clarified lysate (1mg of total protein quantified with BCA assay) was added to streptavidin magnetic beads. The beads were allowed to enrich with the biotinylated proteins overnight at 4°C. After enrichment, beads were washed with RIPA buffer and ice-cold PBS. To remove detergents, the beads were washed with 1 M KCl, 0.1 M Na2CO3, 2 M urea in 10 mM Tris-HCl (pH 8.0), and lastly with 100 mM ammonium hydrogen carbonate (AMBIC). AMBIC was removed, and beads were immediately stored at -80°C until MS processing time.

For WB the biotin labeling was extended to 30 min and the biotinylated proteins were isolated with the same protocol described above. After the protein binding step, the beads were washed with RIPA buffer and PBS, and 2X sample buffer was then added and heated to 95°C for 5 min to elute biotinylated proteins.

### Proteomics analyses

Protein samples were reduced, alkylated, and digested using filter-aided sample preparation (IP samples) or following chloroform/methanol extraction (input lysates) with sequencing grade modified porcine trypsin (Promega). Tryptic peptides were then separated by reverse phase XSelect CSH C18 2.5 um resin (Waters) on an in-line 150 x 0.075 mm column using an UltiMate 3000 RSLCnano system (Thermo Fischer). Peptides were eluted using a 60 min gradient from 98:2 to 65:35 buffer A:B ratio. Eluted peptides were ionized by electrospray (2.2 kV) followed by mass spectrometric analysis on an Orbitrap Exploris 480 mass spectrometer (Thermo). To assemble a chromatogram library, six gas-phase fractions were acquired on the Orbitrap Exploris with 4 m/z DIA spectra (4 m/z precursor isolation windows at 30,000 resolution, normalized AGC target 100%, maximum inject time 66 ms) using a staggered window pattern from narrow mass ranges using optimized window placements. Precursor spectra were acquired after each DIA duty cycle, spanning the m/z range of the gas-phase fraction (i.e. 496-602 m/z, 60,000 resolution, normalized AGC target 100%, maximum injection time 50 ms). For wide-window acquisitions, the Orbitrap Exploris was configured to acquire a precursor scan (385-1015 m/z, 60,000 resolution, normalized AGC target 100%, maximum injection time 50 ms) followed by 50x 12 m/z DIA spectra (12 m/z precursor isolation windows at 15,000 resolution, normalized AGC target 100%, maximum injection time 33 ms) using a staggered window pattern with optimized window placements. Precursor spectra were acquired after each DIA duty cycle.

Buffer A = 0.1% formic acid, 0.5% acetonitrile

Buffer B = 0.1% formic acid, 99.9% acetonitrile

Following data acquisition, data were searched using Spectronaut (Biognosys version 18.3) against the UniProt Homo sapiens database (Proteome ID: UP000005640, Taxon ID: 9606, 3rd version of 2023) using the directDIA method with an identification precursor and protein q-value cutoff of 1%, generate decoys set to true, the protein inference workflow set to maxLFQ, inference algorithm set to IDPicker, quantity level set to MS2, cross-run normalization set to false, and the protein grouping quantification set to median peptide and precursor quantity. Fixed Modifications were set to Carbamidomethyl (C) and variable modifications were set to Acetyl (Protein N-term), Oxidation (M). Protein MS2 intensity values were assessed for quality using ProteiNorm. The data were normalized using Cyclic Loess and analyzed using proteoDA to perform statistical analysis using Linear Models for Microarray Data (limma) with empirical Bayes (eBayes) smoothing to the standard errors. Proteins with an FDR adjusted p-value < 0.05 and a fold change > 2 were considered significant.

Proteomic visualization charts were made using RStudio. All codes have been deposited in GitHub, Lupashin-Lab/wsaragon.

### Western blotting

Protein samples were loaded into the 8-16% Bio-Rad SDS gel and transferred to 0.22 μm nitrocellulose membrane using semi-dry transfer unit (Thermo Fischer). Membranes were stained with Ponceau S stain to assess loading and then blocked in Bio-Rad blocking buffer for subsequent incubation with primary and secondary antibodies (**Table 2**). Washed membranes were scanned on LI-COR Bio Odyssey scanner.

**Table 2.** Antibodies.

| Antibody | Source | Species | WB dilution | IF dilution |
|---|---|---|---|---|
| TurboID | Agrisera, AS20 4440 | Rabbit | 1:2000 | 1:500 |
| LAMP2 | DSHB, H4B4 | Mouse | 1:500 | N/A |
| GPP130 | Covance, PRB-144C | Rabbit | 1:1000 | N/A |
| B4GALT1 | R&D Systems, AF-3609 | Goat | 1:500 | N/A |
| Actin | Sigma, A5441 | Mouse | 1:2000 | N/A |
| VPS50 | Santa Cruz, sc101010 | Mouse | 1:1000 | N/A |
| VPS52 | Bonifacino lab | Rabbit | 1:1000 | N/A |
| VPS53 | Thermo Fisher, PA520548 | Rabbit | 1:1000 | N/A |
| CCDC186 | Invitrogen, PA5-53704 | Rabbit | 1:1000 | 1:400 |
| VPS33B | Protein Tech, 12195-1-AP | Rabbit | 1:2000 | 1:500 |
| GOLGA4/P230 | BD transduction labs. #611280 | Mouse | 1:500 | 1:200 |
| TBC1D23 | Protein Tech, 17002-I-AP | Rabbit | 1:1000 | N/A |
| COG3 | Lupashin lab | Mouse | 1:1000 | N/A |
| TRIP11/GMAP210 | Bethyl, A301-187A | Rabbit | 1:500 | N/A |
| CASP/CUX1 | Thermo Fisher, #PA5-30003 | Rabbit | 1:1000 | N/A |
| Myc tag | Cell Signaling, #2276 | Mouse | N/A | 1:500 |
| TGOLN2/TGN46 | BioRad, AHP500G | Sheep | 1:2000 | 1:500 |
| COG8 | Sigma, SAB4200427 | Rabbit | N/A | 1:500 |
| GALNT2 | R&D Systems, AF7507-SP | Sheep | N/A | 1:500 |
| COPB2 | ABclonal, #A7036 | Rabbit | N/A | 1:400 |
| GOLGA2/GM130 | BD Biosciences, #610823 | Mouse | N/A | 1:500 |
| GOLGA2/GM130 | CalBiochem, #CB1008 | Rabbit | N/A | 1:300 |
| HA tag | Biolegend, #901513 | Mouse | N/A | 1:500 |
| GOLGB1 | Invitrogen, PA5-52772 | Rabbit | N/A | 1:500 |
| WWOX | St. John’s L., STJ114526 | Rabbit | 1:500 | N/A |
| CI-MPR | Abcam, AB124767 | Rabbit | N/A | 1:500 |
| CD-MPR | Abcam, AB134153 | Rabbit | N/A | 1:500 |
| STX7 | R&D, AF5478-SP | Sheep | N/A | 1:200 |
| Fibronectin | Cell Signaling, #26836 | Rabbit | 1:1000 | N/A |
| Cathepsin D | Sigma, C0715 | Mouse | 1:500 | N/A |
| IRDye 800 anti-Mouse | LiCOR/926-68170 | Goat | 1:20000 | N/A |
| IRDye 800 anti-Rabbit | LiCOR/926-32211 | Goat | 1:20000 | N/A |
| IRDye 800 anti-Goat | LiCOR/926-32214 | Donkey | 1:20000 | N/A |
| Alexa Fluor 647 anti-Rabbit | Jackson Immuno Research/711-605-152 | Donkey | 1:2000 | 1:500 |
| Alexa Fluor 647 anti-Mouse | Jackson Immuno Research/715-605-151 | Donkey | 1:2000 | 1:500 |
| Alexa Fluor 647 anti-Goat | Jackson Immuno Research/705-605-147 | Donkey | 1:2000 | 1:500 |
| DyLight 647 anti-Sheep | Jackson Immuno Research/713-605-147 | Donkey | N/A | 1:500 |
| Cy3-anti-Rabbit | Jackson Immuno Research/711-165-152 | Donkey | N/A | 1:500 |
| Cy3-anti-Mouse | Jackson Immuno Research/715-165-151 | Donkey | N/A | 1:500 |
| Alexa Fluor 488 anti-Rabbit | Jackson Immuno Research/711-545-152 | Donkey | N/A | 1:500 |
| Alexa Fluor 488 anti-Mouse | Jackson Immuno Research/715-545-151 | Donkey | N/A | 1:500 |

### Immunofluorescence

Cells were plated on glass coverslips to 80%–90% confluency and fixed with 4% PFA diluted in PBS for 15 min at room temperature. Cells were then permeabilized with 0.1% Triton X-100 in PBS for 1 min, followed by 50 mM ammonium chloride wash for 5 min to remove remaining fixative. Then, cells were blocked with 1% BSA and 0.1% saponin in PBS for 20 min for subsequent incubation with primary antibody (diluted in 1% cold fish gelatin, 0.1% saponin in PBS) for 45 min, washed with PBS and then incubated with fluorescently conjugated secondary antibodies for 30 min and thoroughly washed with PBS to remove unbound antibody. Coverslips were then mounted on glass microscope slides using Prolong Gold antifade reagent (Life Technologies). Cells were imaged with a 63× oil 1.4 numerical aperture objective of an LSM880 Zeiss Laser inverted microscope and Airyscan super-resolution Cocchiaro microscope using ZEN software. Quantitative analysis was performed using single-slice confocal images. All the microscopic images shown are Z-stacked Maximum Intensity Projection images.

### Electron microscopy

Samples were processed using a modified version of the protocol developed by Valdivia and colleagues^89^ with some modifications^90^. Briefly, the excess growth medium was removed from the dish with cells, and an equal volume of 1x fixative was added for 5 min incubation. Then the 0.5x fixative was replaced with 1x fixative for 10 min at RT. The 1x fixative was composed of 4% PFA (EMS) and 1% GA (EMS) in 0.1 M Phosphate buffer, pH 7.4. Finally, cells were fixed on ice for 20 min in 2.5% glutaraldehyde and 0.05% malachite green (EMS) prepared in 0.1 M sodium cacodylate buffer (pH 6.8). Samples were then post-fixed for 30 min at room temperature in 0.5% osmium tetroxide and 0.8% potassium ferricyanide in 0.1 M sodium cacodylate buffer, followed by incubation in 1% tannic acid for 20 min on ice and 1% aqueous uranyl acetate for 1 h at room temperature. Specimens were dehydrated through a graded ethanol series, washed with Propylene Oxide (EMS) and incubated in 50% PO/resin mixture before embedding in Araldite 502/Embed 812 resins (EMS). Ultrathin sections (50 nm) were prepared using a Leica UltraCut UCT ultramicrotome and post-stained with aqueous uranyl acetate and Reynolds’ lead citrate (EMS). Images were acquired on an FEI Tecnai TF20 transmission electron microscope operated at 80 kV and equipped with an FEI Eagle 4k digital camera controlled by FEI acquisition software.

### Golgi localization studies

To form ministacks, cells were treated with 33 μM nocodazole and stained using the IF protocol described. An siRNA-resistant 2×myc-WWOX construct was synthesized by GenScript by introducing synonymous nucleotide substitutions that replaced the siRNA target sequence (5′-CCCATCGATTTACAGATATTA-3′) with 5′-GCCACAGATTCACCGACATCA-3′ without altering the WWOX amino acid sequence. The construct was placed under the control of the COG4 promoter. For the cells transiently expressing myc-WWOX, transfection was done 24 h prior to nocodazole treatment. Location Quotient (LQ) was calculated as described by the Lei Lu lab^91^. Briefly, the distance between the centers of mass of TGOLN2 (*trans*; position 1) and GOLGA2 (*cis*, position 0) was used as the denominator and the distance between the center of mass of Myc-WWOX and GOLGA2 was used as the numerator; the average ratio is the LQ.

### Mitochondria relocalization studies

For mitochondrial relocalization experiments, full-length human CCDC186 and a DNA fragment encoding 3×HA-OMP25^92^ were synthesized by GenScript. To generate an N-terminal mitochondrial targeting construct, CCDC186 was subcloned into a previously described TOM20-Dronpa vector^74^. To generate a C-terminal mitochondrial targeting construct, CCDC186 lacking the stop codon was fused in-frame to 3×HA-OMP25 and cloned into pcDNA3.1. RPE1 cells or HeLa cells were transfected 24 h before IF or EM processing.

### Auxin-inducible degradation assays

Acute inactivation of the COG4 subunit of the COG complex was carried out as previously described^24,31^. Briefly, RPE1 cells stably expressing COG4-HA-mAID and TIR1-V5 under the regulation of the COG4 promoter were treated with 10 µM indole-3-acetic acid (IAA) for 2 h. Degradation efficiency was assessed by IF and WB.

### Transfections

For expression plasmid transfections, cells were grown to 80% confluency and transfected with Lipofectamine 3000 reagents as per manufacturer’s instructions (Thermo Fisher). Cells were incubated in a 37°C incubator with 5% CO2 and 90% humidity for 24 h.

For siRNA KD, cells were grown to 30% confluency and transfected with RNAi MAX lipofectamine reagent as per manufacturer’s instructions (Thermo Fisher). Cells were treated with a final concentration of 100 nM siRNA. WWOX siRNA was published and validated before^67^, target sequence 5′-CCCATCGATTTACAGATATTA-3′. CCDC186 siRNA was purchased from Horizon Discovery, ON-TARGETplus Human CCDC186 (55088) siRNA – SMARTpool. Cells were incubated at the same conditions described above for 72 h.

### Secretion assay

Protocol has been previously described^23^. Briefly, RPE1 cells were grown in complete media and transfected with either control or CCDC186 siRNA for 48 h. After this time, the medium was changed to serum-free media and collected after 36 h. Media was concentrated using 10k concentrator (Amicon Ultra 10k, Millipore) to 20X. For WB, secretion was normalized to the number of cells in the total lysate.

---

**Supplementary Figure 1. Validation of TurboID fusion protein localization and VPS53-dependent proximity labeling.** (**A**) IF analysis of RPE1 cells stably expressing GFP-TurboID as a proximity-labeling control. GFP-TurboID exhibits diffuse intracellular distribution and widespread protein biotinylation, detected using fluorescent streptavidin. GOLGA4 marks the Golgi apparatus. Individual fluorescence channels and merged images are shown. Scale bars, 10 µm. (**B-D**) IF analysis of RPE1 cells stably expressing VPS50-TurboID (**B**), VPS52-TurboID (**C**), or VPS54-TurboID (**D**) following transient expression of mGold-RAB4A. VPS50-TurboID and VPS52-TurboID (red) colocalize with mGold-RAB4A-positive endosomes (green), whereas VPS54-TurboID remains predominantly in the perinuclear Golgi region and shows little overlap with RAB4A-positive compartments. These results support the association of VPS50 and VPS52 with EARP-positive endosomes and the preferential Golgi localization of the GARP-specific subunit VPS54. (**E**) IF analysis of HA-VPS33B and mGold-RAB4A in RPE1 cells. VPS33B partially colocalizes with RAB4A-positive endosomes, identifying a potential compartment for functional cooperation between EARP and the VPS33B–VIPAS39 (CHEVI) complex. Boxed regions are enlarged on the right, with individual channels displayed in grayscale. Scale bars, 10 µm.

**Supplementary Figure 2. Selective proximity labeling of Golgi and endolysosomal multisubunit complexes by CATCHR-TurboID constructs.** Dot plots showing the relative abundance and enrichment of subunits of Golgi and endolysosomal multisubunit complexes identified by TurboID proximity labeling. Dot size represents average protein abundance, and color indicates log₂ fold enrichment relative to GFP-TurboID controls. (**A**) Proximity labeling of CATCHR complex subunits. VPS50-, VPS52-, and VPS53-TurboID preferentially label EARP subunits, whereas VPS52-, VPS53-, and VPS54-TurboID label GARP subunits. VPS54-TurboID also labels several COG subunits. Conversely, COG4- and COG6-TurboID label all eight COG subunits, as well as the GARP subunits VPS53 and VPS54. These overlapping labeling patterns reveal distinct but partially shared molecular environments of Golgi and endosomal CATCHR complexes. (**B**) Proximity labeling of endolysosomal tethering complex subunits. EARP-associated TurboID constructs preferentially label VPS33B and VIPAS39, the two subunits of the CHEVI complex, whereas subunits of the CORVET, HOPS, and FERARI complexes show little or no enrichment. These results support a selective spatial association between EARP and CHEVI within the endosomal trafficking system.

**Supplementary Figure 3. CCDC186 localizes to the *trans*-Golgi region and mediates membrane tethering through its C-terminal region.** (**A**) High-resolution immunofluorescence analysis of endogenous CCDC186 (red) localization relative to the trans-Golgi marker TGOLN2/TGN46 (blue) and the cis-Golgi marker GOLGA2/GM130 (green) in RPE1 cells. Cells were treated with nocodazole to disperse the Golgi ribbon into individual ministacks, allowing more precise analysis of CCDC186 localization within the Golgi. The fluorescence intensity profile along the indicated line demonstrates that CCDC186 localizes predominantly to the trans-Golgi region, closely associated with TGOLN2/TGN46. Four additional representative Golgi ministacks (R2–R5) are shown. (**B-C**) Analysis of CCDC186-mediated membrane tethering following its ectopic targeting to mitochondria. (**B**) IF analysis of RPE1 cells transiently expressing CCDC186 anchored to the outer mitochondrial membrane through either its N-terminus (TOM20-CCDC186) or C-terminus (CCDC186-OMP). Both constructs alter mitochondrial network organization. (**C**) TEM of cells expressing CCDC186-OMP reveals electron-dense connections between adjacent mitochondrial membranes (arrowheads). In contrast, N-terminally anchored CCDC186 promotes vesicle accumulation and tethering at mitochondria, as shown in Figure 5B. These observations suggest that the C-terminal region of CCDC186 contributes to vesicle tethering. (**D**) Quantification of the colocalization between GOLGA4/P230 and mitochondria expressing control or mitochondrially targeted CCDC186 constructs. Pearson’s colocalization coefficients show no significant difference between the two conditions (ns, not significant). (**E-F**) Validation of siRNA-mediated CCDC186 depletion in RPE1 cells. (**E**) Representative immunoblot showing reduced endogenous CCDC186 levels following transfection with pooled CCDC186-targeting siRNAs. β-Actin serves as a loading control. (**F**) Quantification of CCDC186 protein levels normalized to β-actin, confirming significant depletion compared with control cells (*P* < 0.05). (**G**) Immunoblot analysis of total cell lysates (lanes 1–4) and concentrated conditioned media (lanes 5–8) from wild-type, CCDC186 KD, VPS50 KO, and VPS54 KO RPE1 cells. CCDC186 deficiency alters extracellular fibronectin abundance without producing the pronounced increase in secreted pro-cathepsin D observed in VPS54 KO cells. β-Actin serves as a loading control for cell lysates.

**Supplementary Figure 4. WWOX localizes to the medial Golgi and contributes to Golgi organization and glycosylation.** (**A-B**) High-resolution immunofluorescence analysis of myc-WWOX (red) localization relative to the COG complex subunit COG8 (green; **A**) and the Golgi matrix protein GOLGB1/giantin (green; **B**). RPE1 cells were treated with nocodazole to disperse the Golgi ribbon into individual ministacks, allowing more precise analysis of WWOX localization within the Golgi. myc-WWOX exhibits a ring-like distribution closely associated with COG8- and giantin-positive Golgi membranes. Representative Golgi ministack and corresponding fluorescence intensity profile along the indicated line is shown, together with four additional ministacks (R2–R5). (**C**) Immunoblot analysis confirming efficient siRNA-mediated depletion of WWOX in HEK293 cells. β-Actin serves as a loading control. (**D**) Representative IF images of control (CTRL) and WWOX KD RPE1 cells stained for the cis-Golgi marker GOLGA2. WWOX depletion alters Golgi morphology, consistent with a role for WWOX in maintaining normal Golgi organization. Scale bars, 50 µm. (**E**) Pearson’s colocalization coefficient of CTRL and WWOX KD in RPE1 cells. (**F**) Representative immunofluorescence images of WWOX-depleted HEK293 cells expressing myc-tagged WWOX, stained with HPA (blue), anti-myc (red), and anti-GOLGA2 (green). Expression of siRNA-resistant myc-WWOX suppresses the increased HPA staining associated with WWOX depletion, indicating rescue of the O-glycosylation defect. White arrowheads indicate cells expressing myc**-**WWOX and exhibiting reduced HPA staining and grey arrowheads indicate cells not expressing myc-WWOX. Scale bars, 10 µm.

## Acknowledgements

We thank our colleagues who generously provided reagents used in this study. We are particularly grateful to Tetyana Kudlyk for technical assistance with the generation of CATCHR-TurboID constructs and to the members of the Lupashin laboratory for valuable discussions and critical feedback. We also thank the UAMS Digital Microscopy, Proteomics, and DNA Sequencing Cores for access to their facilities and for their expert technical support.

## Abbreviations

CATCHR: Complexes Associated with Tethering Containing Helical Rods
CCT: Coiled-Coil Tether
COG: Conserved Oligomeric Golgi
GARP: Golgi-Associated Retrograde Protein
EARP: Endosome-Associated Recycling Protein
IF: Immunofluorescence
KD: Knock Down
KO: Knock Out
MS: Mass Spectrometry
NRZ: NAG, RINT1, and ZW10
SM: Sec1/Munc18
SNARE: Soluble N-ethylmaleimide-sensitive factor Attachment protein Receptor
TEM: Transmission Electron Microscopy
WB: Western blot

