## Supplementary Figures 1-4 for "Golgi-endosomal CATCHR complexes function as organizing hubs for vesicle tethering and fusion"

**Supplementary figure 1:** Validation of TurboID fusion protein localization and VPS53-dependent proximity labeling.

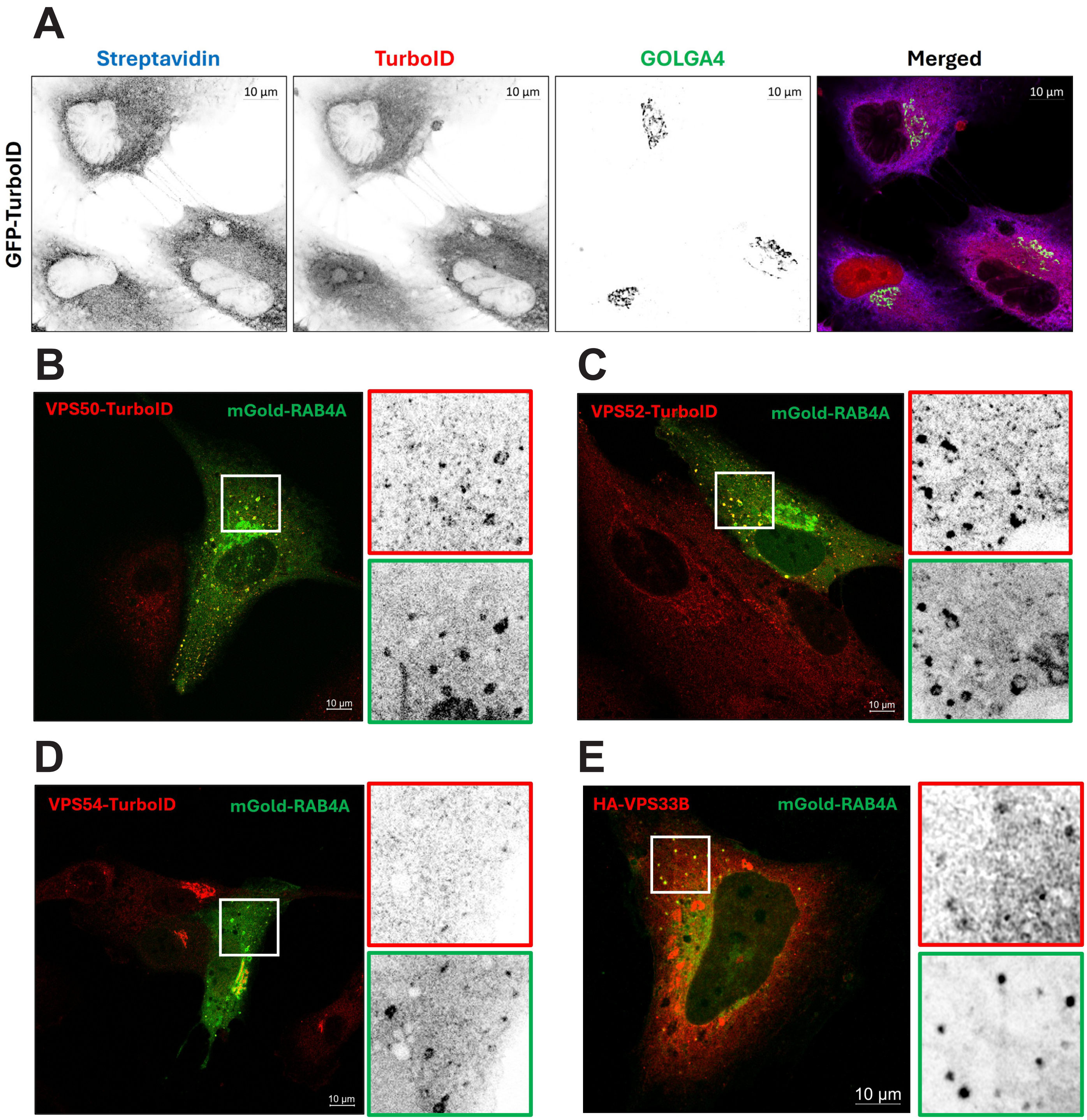

**Supplementary figure 2:** Selective proximity labeling of Golgi and endolysosomal multisubunit complexes by CATCHR-TurboID constructs.

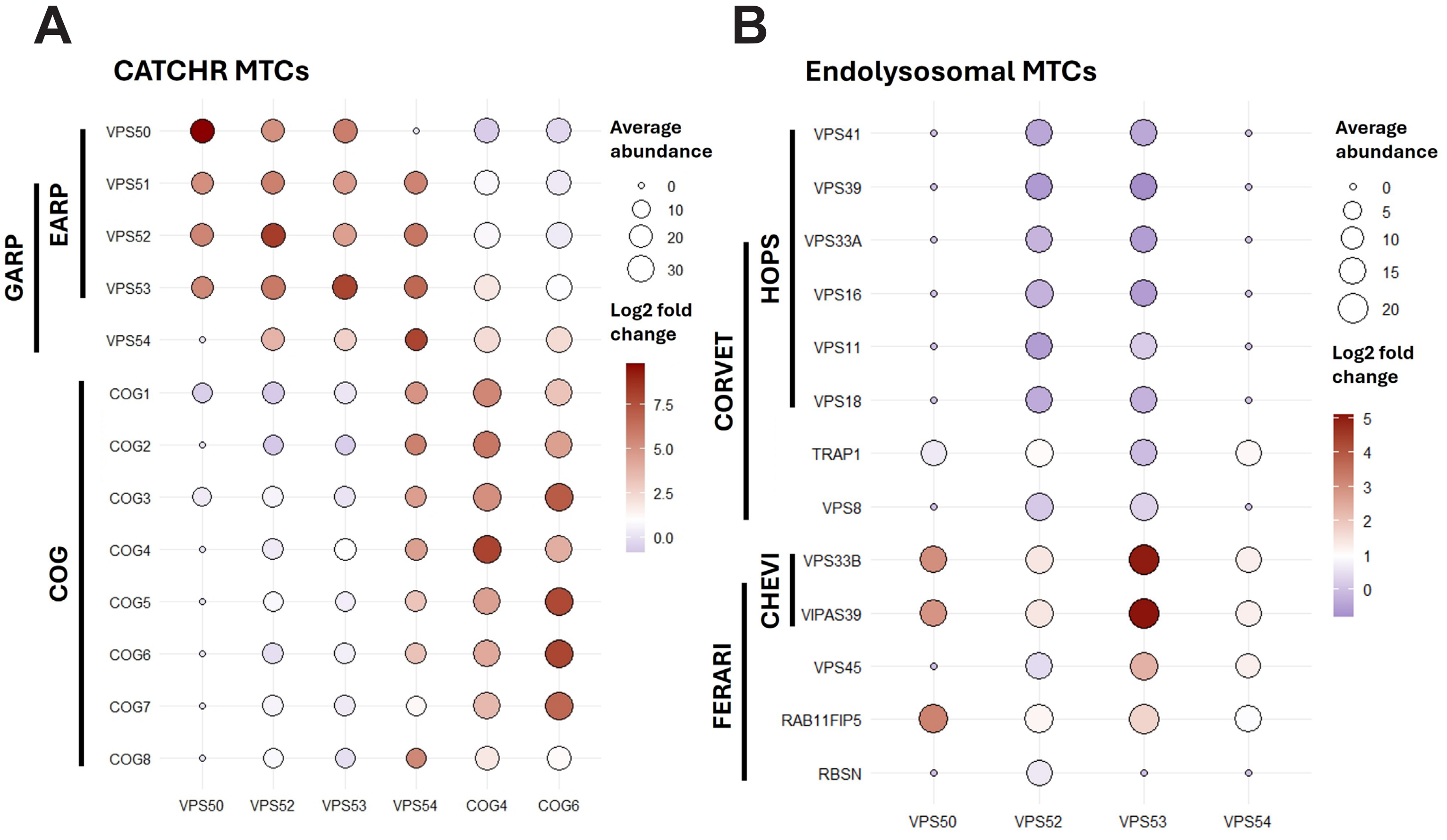

**Supplementary figure 3:** CCDC186 localizes to the trans-Golgi region and mediates membrane tethering through its C-terminal region.

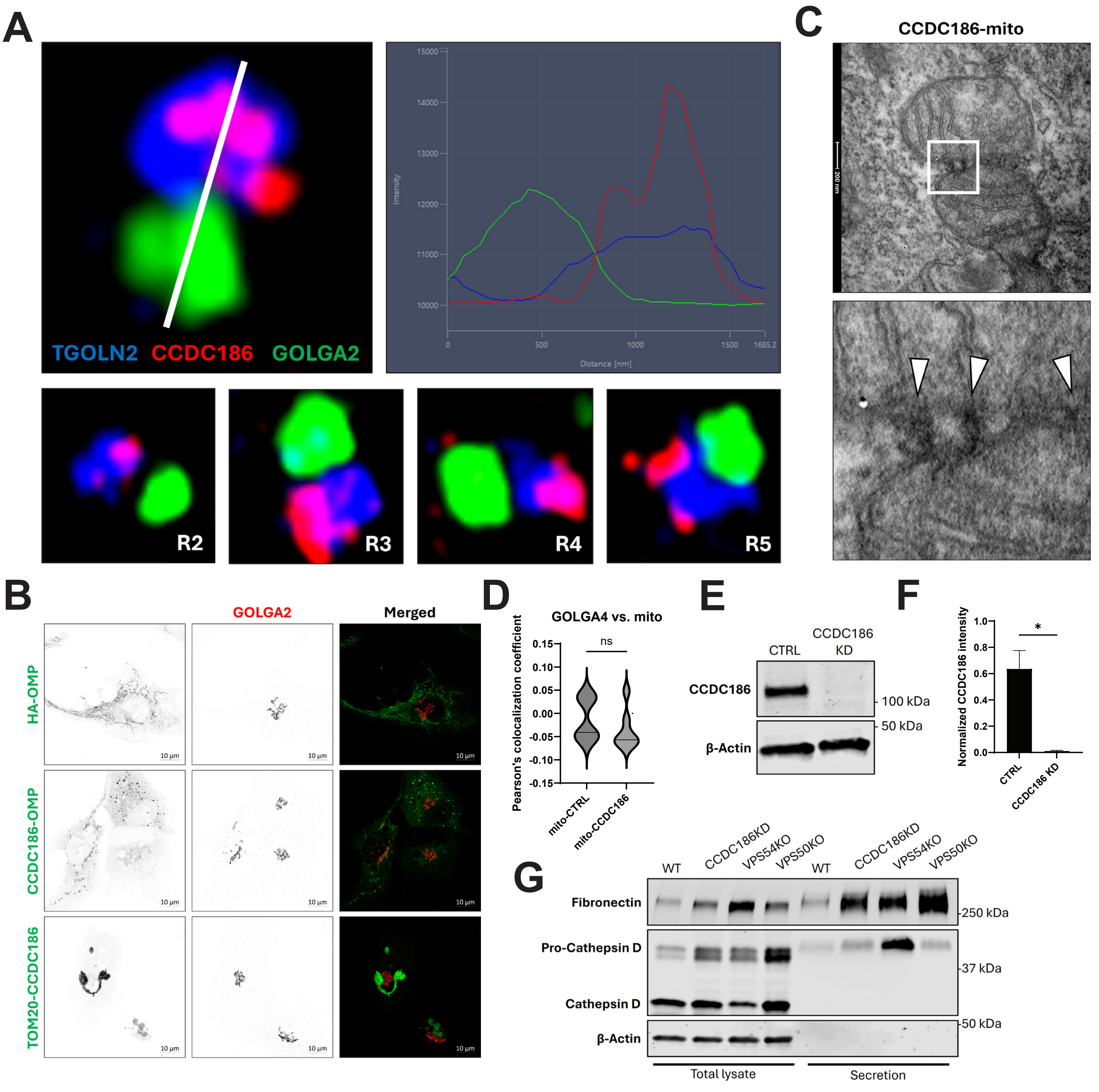

**Supplementary figure 4:** WWOX localizes to the medial Golgi and contributes to Golgi organization and glycosylation.

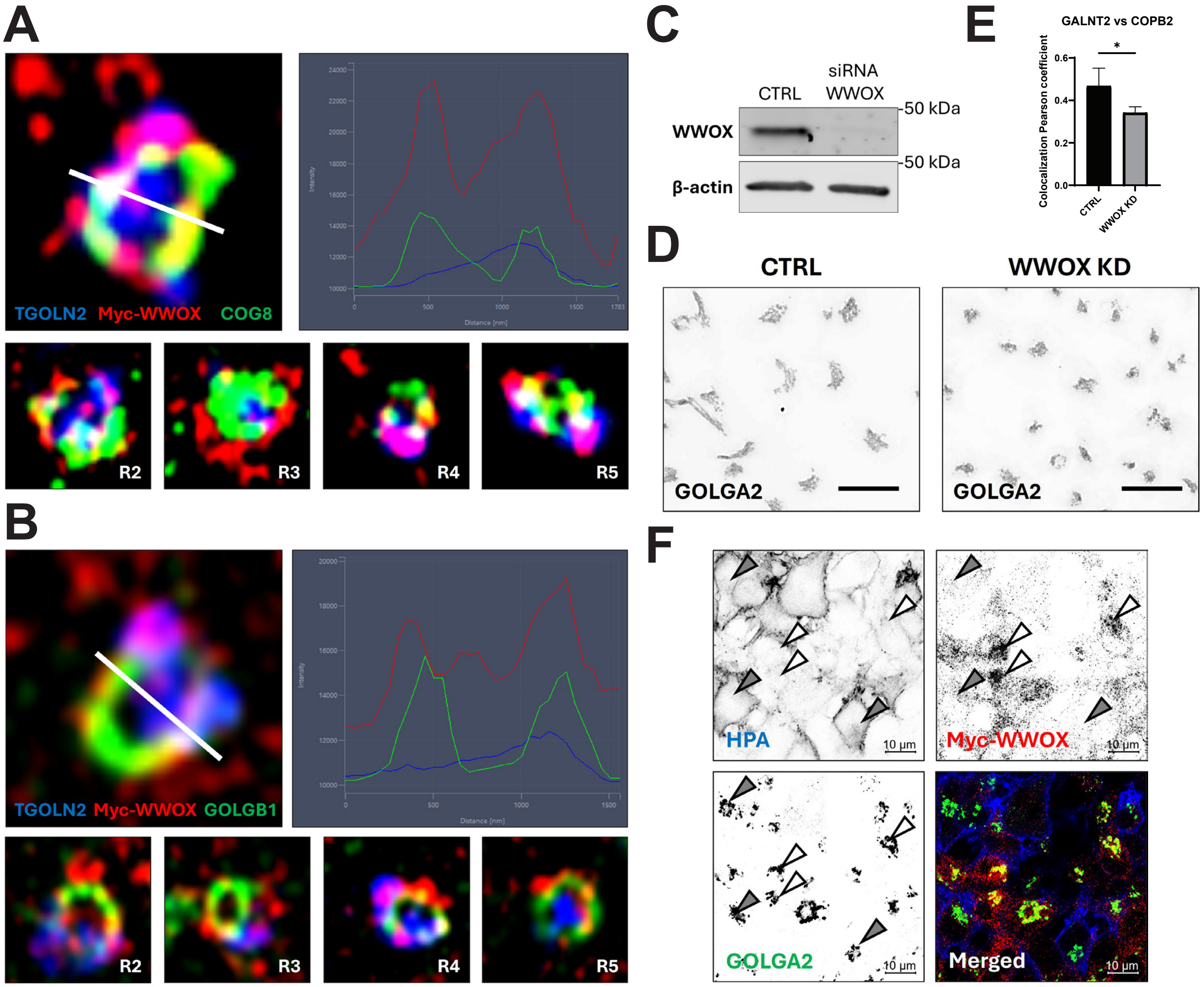
